# Comprehensive phenotypic analysis of the Dp1Tyb mouse strain reveals a broad range of Down Syndrome-related phenotypes

**DOI:** 10.1101/2021.02.11.430828

**Authors:** Eva Lana-Elola, Heather Cater, Sheona Watson-Scales, Simon Greenaway, Jennifer Müller-Winkler, Dorota Gibbins, Mihaela Nemes, Amy Slender, Tertius Hough, Piia Keskivali-Bond, Cheryl L Scudamore, Eleanor Herbert, Gareth T Banks, Helene Mobbs, Tara Canonica, Justin Tosh, Suzanna Noy, Miriam Llorian, Patrick M. Nolan, Julian L. Griffin, Mark Good, Michelle Simon, Ann-Marie Mallon, Sara Wells, Elizabeth M. C. Fisher, Victor L. J. Tybulewicz

## Abstract

Down syndrome (DS), trisomy 21, results in many complex phenotypes including cognitive deficits, heart defects and craniofacial alterations. Phenotypes arise from an extra copy of human chromosome 21 (Hsa21) genes. However, causative genes remain mostly unknown. Animal models enable identification of these genes and pathological mechanisms. The Dp1Tyb mouse model of DS has an extra copy of 63% of Hsa21-orthologous mouse genes. Here, we comprehensively phenotype Dp1Tyb mice and find wide-ranging DS-like phenotypes including aberrant megakaryopoiesis, reduced bone density, and deficits in memory, locomotion, hearing and sleep. Thus, Dp1Tyb mice are an excellent model for studies of many complex DS phenotypes.

## Introduction

Down syndrome, which arises from trisomy of human chromosome 21 (Hsa21), is a complex condition comprising of a large number of phenotypes, which differ in severity and penetrance^1^. Individuals with DS have learning and memory deficits, shorter stature, reduced bone density, craniofacial alterations and increased frequencies of congenital heart defects, leukaemia, diabetes, motor deficits, disrupted sleep, and impaired hearing and vision. People with DS are high-risk for Alzheimer’s disease (AD); ∼50% have clinical signs of dementia by age 60^2^. With a prevalence of ∼1 in 800 live births, DS is the most common genetic cause of intellectual disability and of AD^2, 3^.

Although trisomy 21 has been known as the cause of DS since 1959^4, 5^, there are still no effective treatments for most DS phenotypes, in particular the cognitive aspects. DS phenotypes most likely result from increased dosage of genes on Hsa21, which comprise ∼230 coding genes, and many more non-coding elements^3^. Identification of dosage-sensitive Hsa21 genes that cause DS phenotypes would facilitate studies of the underlying pathomechanisms, and development of new treatments. However, for most DS phenotypes these causative genes are unknown^6^.

A promising approach to the identification of these genes is to model the syndrome in mice, and then use the power of mouse genetics to investigate pathology. Two approaches have been taken. Firstly, generating mice with an extra human chromosome comprising most of Hsa21, namely the Tc1 and TcMac21 mouse strains^7, 8^. Secondly, using chromosome engineering to create mouse strains with an additional copy of regions orthologous to Hsa21^9, 10^. The largest region of Hsa21 orthology (23Mb) is found on mouse chromosome 16 (Mmu16), with smaller regions on Mmu10 (3Mb) and Mmu17 (2Mb)^9^. The Dp1Tyb and Dp1Yey strains have an extra copy of the entire Hsa21-orthologous region of Mmu16^11, 12^. Many further strains have been generated with an extra copy of smaller Hsa21-orthologous regions, allowing mapping of causative genes^11, 13–20^.

While various DS mouse models have been investigated for individual specific phenotypes, there has been no comprehensive analysis to ascertain if a single DS model recapitulates the breadth of DS phenotypes. This is a critical issue because some phenotypes likely arise from interactions of linked genes in DS and no ‘chromosomal’ mouse models have yet been validated across a wide-range of phenotypes, relating them to the human condition. Here we use the extensive range of tests in the International Mouse Phenotyping Consortium (IMPC)^21^, augmented with a series of further bespoke assays to carry out a broad phenotypic analysis of the Dp1Tyb mouse strain. This strain has an extra copy of 148 coding genes on Mmu16, comprising 63% of Hsa21-orthologous genes in the mouse genome, and thus replicates the majority of the gene dosage increase in DS, making it an excellent genetic model^10, 11^. We find that Dp1Tyb mice have many phenotypes similar to those seen in people with DS, including decreased bone density, craniofacial changes, altered cardiac function, aberrant erythropoiesis and megakaryopoiesis, a pre-diabetic state, defective hearing, learning and memory deficits, impaired motor activity, and disrupted sleep. Thus, Dp1Tyb mice faithfully recapitulate complex and wide-ranging phenotypes found in human DS and are an excellent tool for discovering causative genes and their pathological mechanisms, including additive and interactive gene effects.

## Results

### Phenotyping pipelines

To carry out a broad phenotypic analysis of Dp1Tyb mice, we bred multiple cohorts of mice and analysed them using a variety of procedures, many arranged into pipelines, with animals given specific tests at particular ages (Fig. 1). We used the IMPC phenotyping pipeline^21^ to study a cohort of 10 female and 10 male Dp1Tyb hemizygous mice and similar numbers of control wild-type (WT) animals produced from a Dp1Tyb x C57BL/6J cross; all animals were congenic on C57BL/6J (cohort 1). This pipeline covers a wide range of physiological systems (Fig. 1 and Methods). Since cognitive and behavioural changes are DS hallmarks, we set up a second pipeline for neurological aspects using 15 male and 15 female mice per genotype (cohorts 2 and 3) and extended it with one further test (cohort 6, object-in-place test). To investigate the haematopoietic system, we carried out flow cytometric analysis (cohort 4). Finally, we analysed older mice ∼1 year of age, for pathological changes and cardiac function (cohort 5).

**Figure 1.**
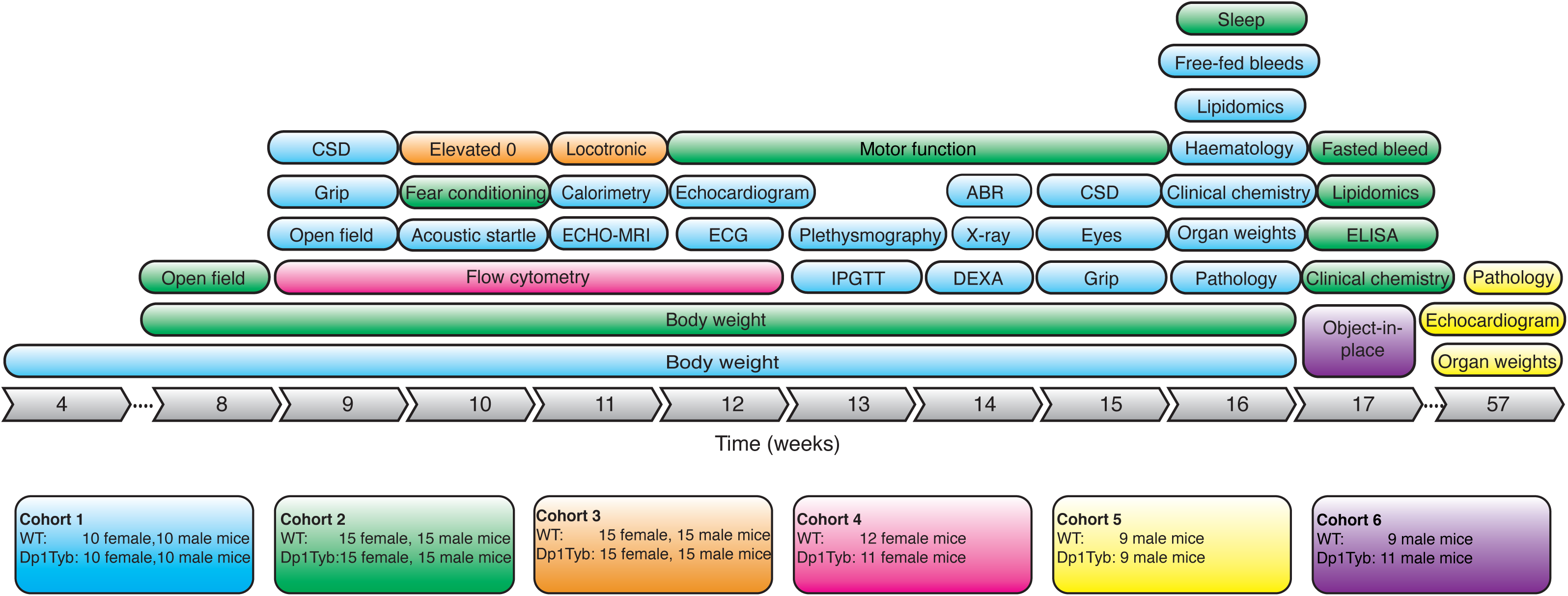
Broad phenotyping pipeline used for analysis of Dp1Tyb mice. Diagram shows the 6 cohorts of Dp1Tyb mice and WT controls that were used in the broad phenotyping analysis. Each cohort is indicated in a different colour, showing the tests it underwent, the numbers and sex of mice involved and the age at which the tests were administered. CSD, Combined SHIRPA and Dysmorphology test.; ECG, electrocardiogram; Elevated 0, elevated 0 maze; IPGTT, intra-peritoneal glucose tolerance test; ABR, acoustic brainstem response; DEXA, dual energy X-ray absorption; ELISA, enzyme-linked immunosorbent assay for plasma hormones.

In total, mice were analysed in 28 different procedures, generating data for 1800 parameters (where both sexes were used, males, females and both sexes together are counted as separate parameters). Statistical analysis showed that using a false discovery rate of less than 5% (q≤0.05), Dp1Tyb mice had significant changes in 468 parameters from 22 procedures compared to WT mice (**Supplementary Table 1**). Generally, where there were differences between Dp1Tyb and WT mice, these were similar for both sexes, with only 15 parameters in 6 procedures showing significant sexual dimorphism (q≤0.05). It has been proposed that aneuploidy results in greater phenotypic variability^22^. To assess this in Dp1Tyb mice, we investigated if the variance of the 1450 parameters that had a numerical value was significantly different (q≤0.05) between the two genotypes. We found that just 39 parameters showed a significant difference in variance, with 22 and 17 of these showing larger variation in WT and Dp1Tyb mice, respectively. Furthermore, there was no significant difference in the coefficient of variation for these 1450 parameters between either female or male Dp1Tyb mice compared to WT controls (Supplementary Fig. 1a, **Supplementary Table 1**). Thus, Dp1Tyb mice do not show greater phenotypic variation in the parameters measured in this study.

### Decreased viability of Dp1Tyb mice

Analysis of pups at weaning from the Dp1Tyb x C57BL/6J cross, showed a very significant decrease in the expected (50%) proportion of female and male Dp1Tyb animals (Supplementary Fig. 1b); 46% and 61% of expected female and male Dp1Tyb pups survived to weaning, respectively. This loss most likely occurs around birth with mothers eating pups, since we did not observe dead pups in the cages, and the proportion of Dp1Tyb embryos was not altered at embryonic day 14.5 (E14.5) of gestation^11^.

### Increased expression of duplicated genes in Dp1Tyb hippocampus

We investigated if the additional copy of 23Mb of Mmu16 in Dp1Tyb mice results in increased expression of the 148 coding genes contained in this region. RNA sequencing of hippocampus from adult Dp1Tyb and WT mice showed that of the 87 expressed genes in the duplicated region, 75 were significantly upregulated, and no genes were significantly downregulated (Fig. 2). The mean(±SD) upregulation of these genes was 1.43(±0.28)-fold, similar to the expected 1.5-fold increase. Thus, the additional 23Mb region of Mmu16 results in increased gene expression in line with gene copy number.

**Figure 2.**
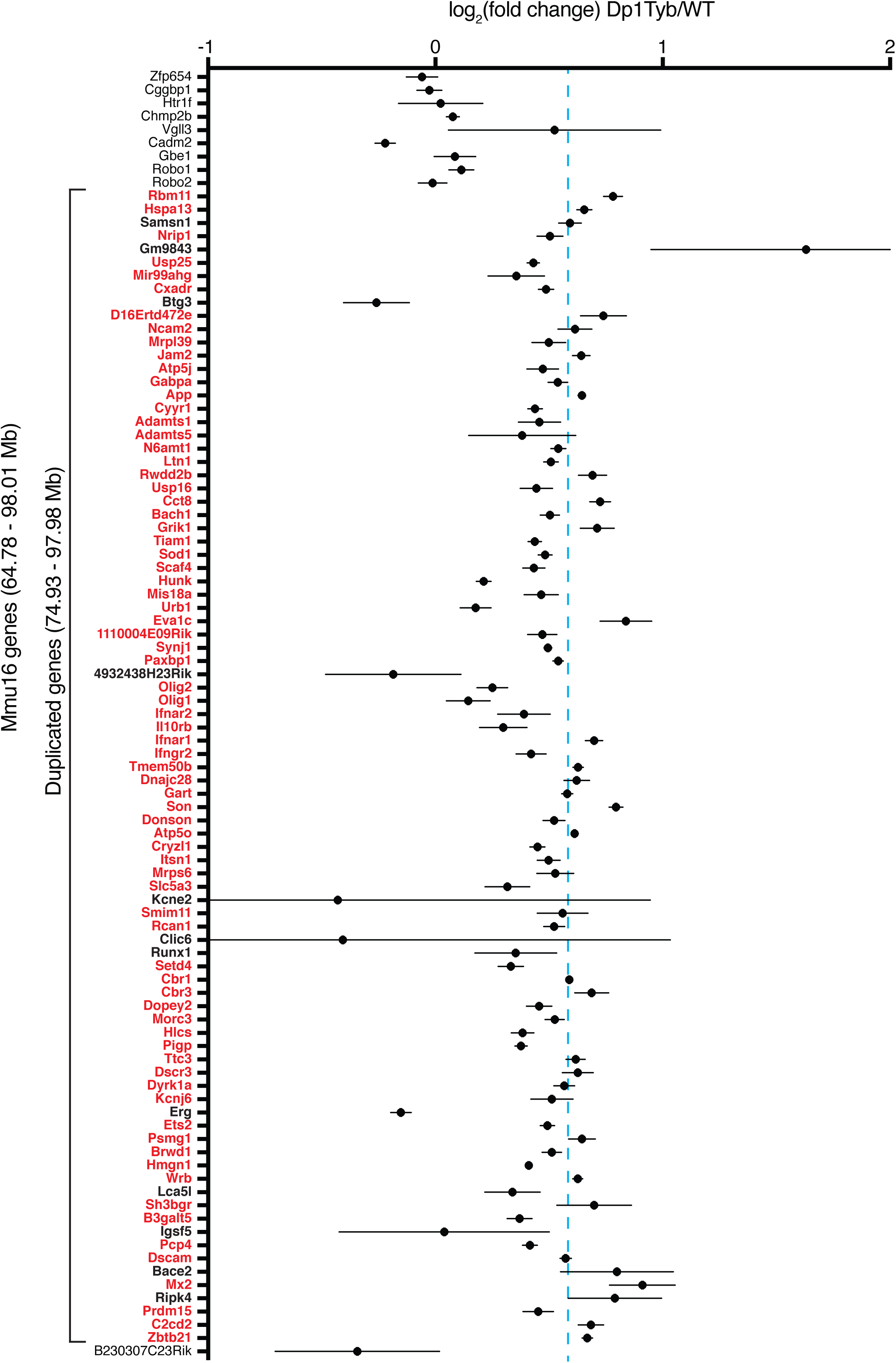
Upregulation of expression of genes in duplicated region of Dp1Tyb mice. Mean±SEM log_2_ fold change of gene expression in the hippocampus between Dp1Tyb and WT mice. Only expressed coding genes and one miRNA host gene shown; expressed genes were defined by sum of expression over all 10 samples > 1 TPM and having a measured p-value in DEseq2. Genes within the duplicated region of Mmu16 are shown in bold. Genes showing a significantly different expression in Dp1Tyb compared to WT (adjusted p-value <0.05) are listed in red. Ten genes that are not duplicated (9 on the centromeric and 1 on the telomeric side of the duplication) are included in the figure to allow comparison with duplicated genes. Dashed blue line indicates a fold change of 1.5 expected by the increased dosage of the duplicated genes.

### Dp1Tyb mice have altered skeletal development

Body weights from 4 to 16 weeks of age, showed that male Dp1Tyb mice weighed less than WT mice at 4 weeks, but caught up afterwards, with no differences in adult mice (Fig. 3a).

**Figure 3.**
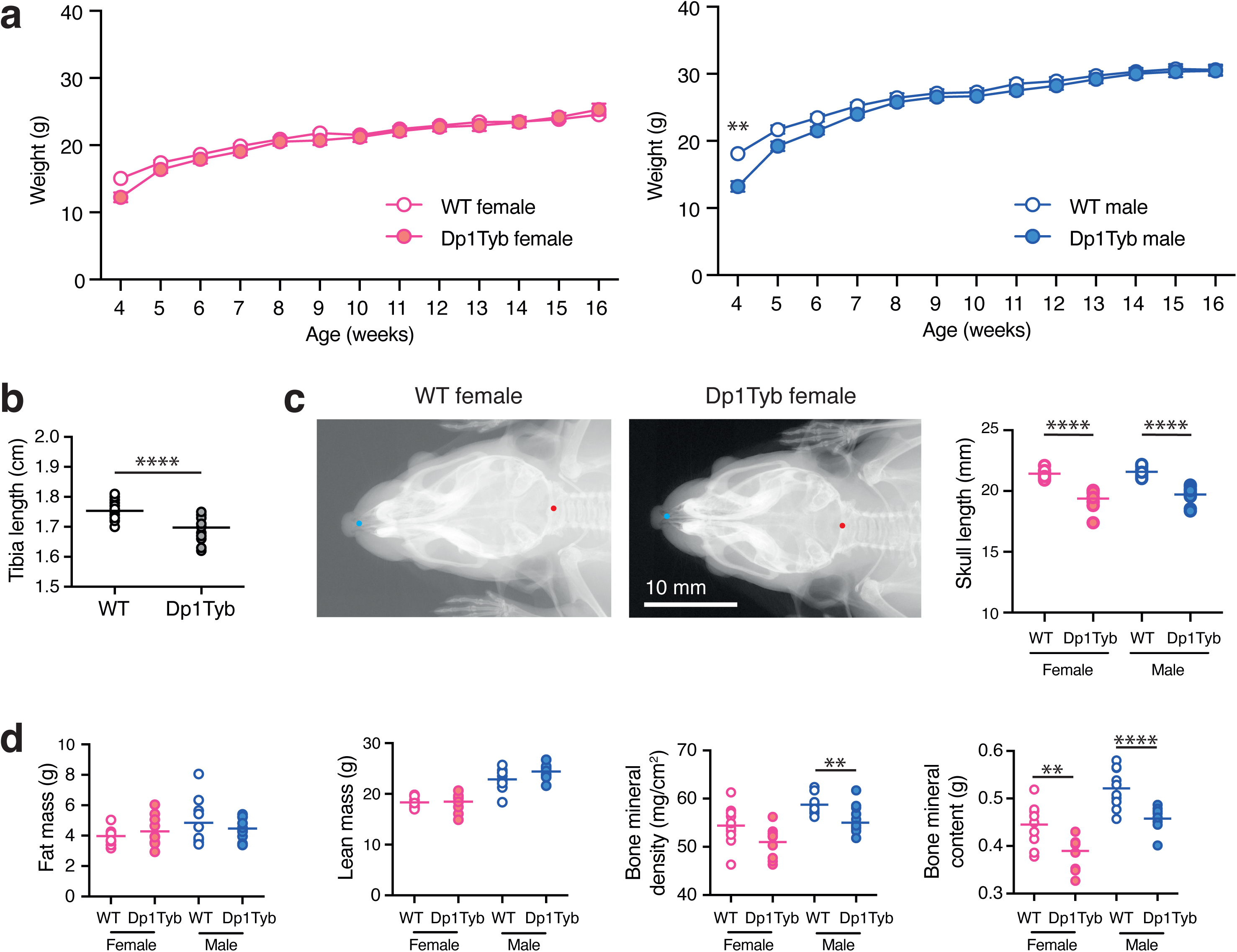
Skeletal changes and decreased bone mineral density and content in Dp1Tyb mice. **a**, Mean±SEM weight of Dp1Tyb and WT mice (cohort 1) as a function of age. **b**, Tibia length of WT and Dp1Tyb mice (cohort 1) showing combined data from females and males, determined from X-ray images. **c**, Example X-ray images of dorsal views of WT and Dp1Tyb female mouse skulls (cohort 1). Graph shows the length of the skull from the anterior tip of the nasal bone to the posterior of the occipital bone, which are indicated in blue and red dots on the images, from WT and Dp1Tyb mice (cohort 1). **d**, Fat mass, lean mass, bone mineral density and bone mineral content of Dp1Tyb and WT mice (cohort 1) determined by DEXA. Horizontal lines indicate mean. Here and in other figures where sexes are analysed separately, the graphs are coloured red and blue for females and males, respectively. Where both sexes are analysed together graphs use white and grey to distinguish the genotypes. ** 0.001 < *q* < 0.01; **** *q* < 0.0001.

People with DS are typically shorter in stature, have lower bone density and characteristic changes in craniofacial morphology, such as brachycephaly (front to back shortening of the skull), and increased body fat content^23–28^. X-ray analysis at 14 weeks of age showed that Dp1Tyb mice had shorter tibia and skulls (Fig. 3b, c), and the Combined SHIRPA and Dysmorphology (CSD) analysis^29^ revealed that Dp1Tyb mice had abnormally shaped heads, snouts and lips (Supplementary Tables 2, 3). Dual Energy X-Ray Absorption (DEXA) showed that Dp1Tyb mice had reduced bone mineral density and content, whereas there was no change in fat or lean mass as measured by DEXA or ECHO-MRI (Fig. 3d, Supplementary Fig. 1c). Thus, Dp1Tyb mice have substantially altered bone growth resulting in skeletal dysmorphology.

### Dp1Tyb mice have enlarged spleens but no Aβ deposition in the hippocampus

Analysis of organ weights showed that at 16 weeks Dp1Tyb mice had slightly altered weights of heart and kidneys and at 57 weeks had reduced liver weights (Supplementary Fig. 1d, e). However, the largest change was an increase in the weights of Dp1Tyb spleens at both ages (Supplementary Fig. 1d, e). Histology showed increased extramedullary haematopoiesis in Dp1Tyb spleens (Supplementary Fig. 2a, Supplementary Table 4). Significant changes were also seen in the portal areas of the liver with bile duct hyperplasia and vascular anomalies (Supplementary Fig. 2b). Otitis media was also observed in all Dp1Tyb animals.

Histological analysis of other organs in Dp1Tyb mice showed no significant findings with any changes recorded being within the expected normal variation in background pathology for the wild type mouse strain. Given the high prevalence of AD in DS^2^, we also examined the hippocampus of 1-year old mice for the presence of Aβ amyloid. While Aβ deposition was readily seen in the J20 transgenic overexpression mouse model of AD, we detected no deposition in Dp1Tyb or WT control mice (Supplementary Fig. 2c).

### Increased metabolic rate in Dp1Tyb mice

People with DS have a reduced resting metabolic rate^30, 31^. In contrast, indirect calorimetry showed that Dp1Tyb mice produced more CO_2_ (VCO_2_), used more O_2_ (VO_2_), and had higher heat production indicating that Dp1Tyb mice have an increased metabolic rate and had an elevated respiratory exchange ratio (RER) (Supplementary Fig. 3a). The latter suggests that the mice may be using a higher ratio of carbohydrate to fat as a fuel source.

### Dp1Tyb mice have altered heart function

∼40% of neonates with DS have congenital heart defects, typically ventricular or atrioventricular septal defects (VSD or AVSD)^32^. We found that Dp1Tyb embryos have a high prevalence of VSD and AVSD at E14.5 of gestation, resembling the defects seen in DS^11^. Analysis of heart function at 12 weeks by echocardiography showed that compared to WT mice, Dp1Tyb animals had a slower heart rate, increased stroke volume, increased cardiac output, increased end-diastolic and end-systolic diameters, and increased left ventricular internal diameter at diastole and systole (Fig. 4a). ECG measurements showed longer QT, corrected QT (QTc), JT and T peak times and larger amplitude of the R wave, consistent with increased left ventricle size (Fig. 4b, c). These changes were not progressive, since echocardiography showed no changes in cardiac function in 57-week old Dp1Tyb mice (Supplementary Fig. 3b).

**Figure 4.**
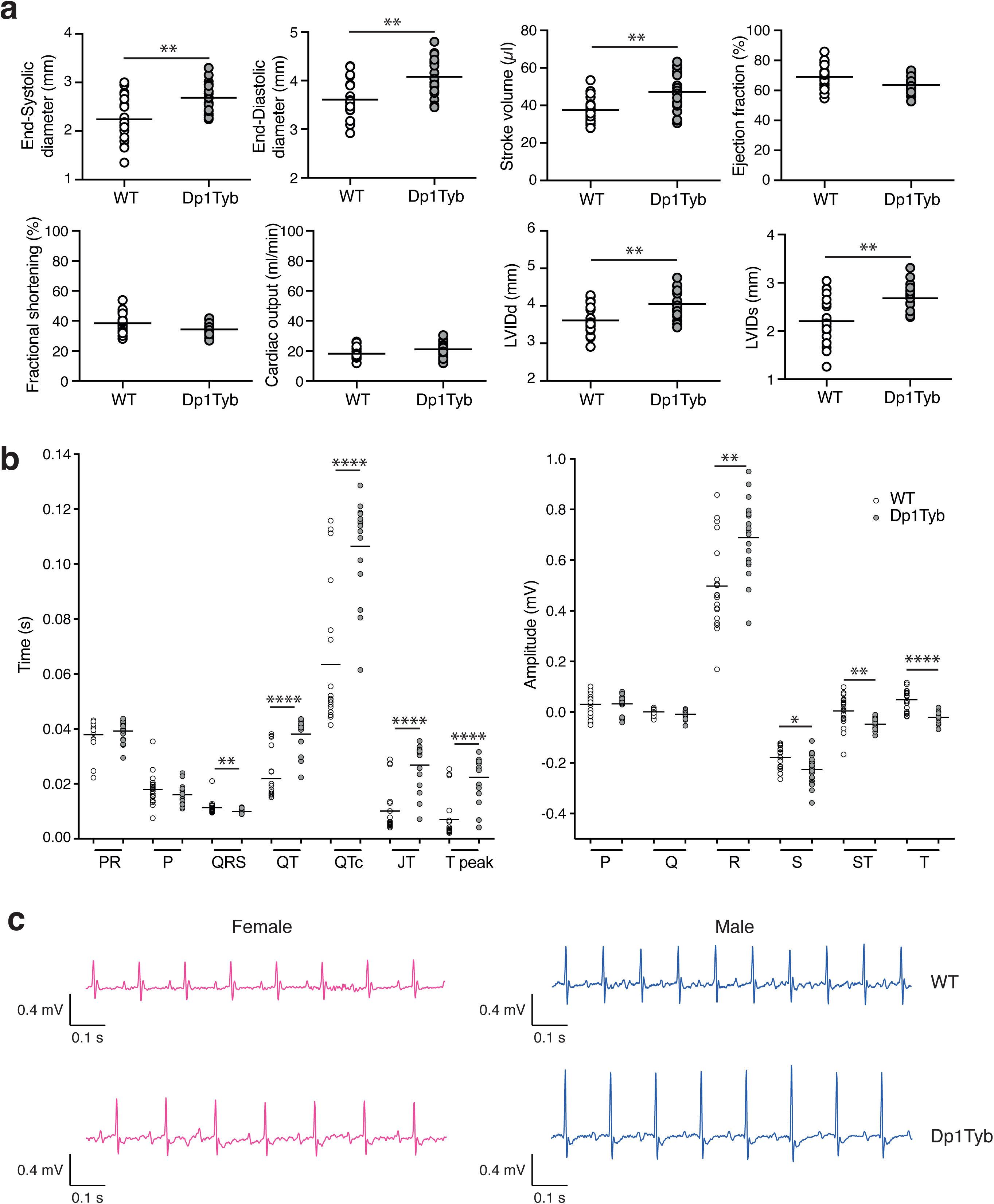
Altered cardiac function in Dp1Tyb mice. **a**, End-systolic diameter, end-diastolic diameter, stroke volume, ejection fraction, fractional shortening, cardiac output, left ventricular inner diameter in diastole (LVIDd) and left ventricular inner diameter in systole (LVIDs) in WT and Dp1Tyb mice (females and males combined) from cohort 1 determined by echocardiography. **b**, mean interval durations and wave amplitudes from electrocardiogram (ECG) analysis of WT and Dp1Tyb mice (females and males combined) from cohort 1. **c**, Representative ECG traces. Horizontal lines indicate mean. * 0.01 < *q* < 0.05; ** 0.001 < *q* < 0.01; **** *q* < 0.0001.

### Increased breathing volumes in Dp1Tyb mice

To analyse lung function, mice were placed into whole body plethysmography chambers and breathing recorded for 5 min in normoxic conditions, then for 5 min in hypoxia (10% O_2_/3% CO_2_) and finally the mice were returned to normoxia for 5 min. WT and Dp1Tyb mice responded to hypoxia by increasing breathing volumes as measured per minute and per breath (tidal). However, under both normoxic and hypoxic conditions, Dp1Tyb mice had increased minute and tidal volumes compared to WT mice (Fig. 5a). This increased rate of breathing may be related to the increased consumption of O_2_.

**Figure 5.**
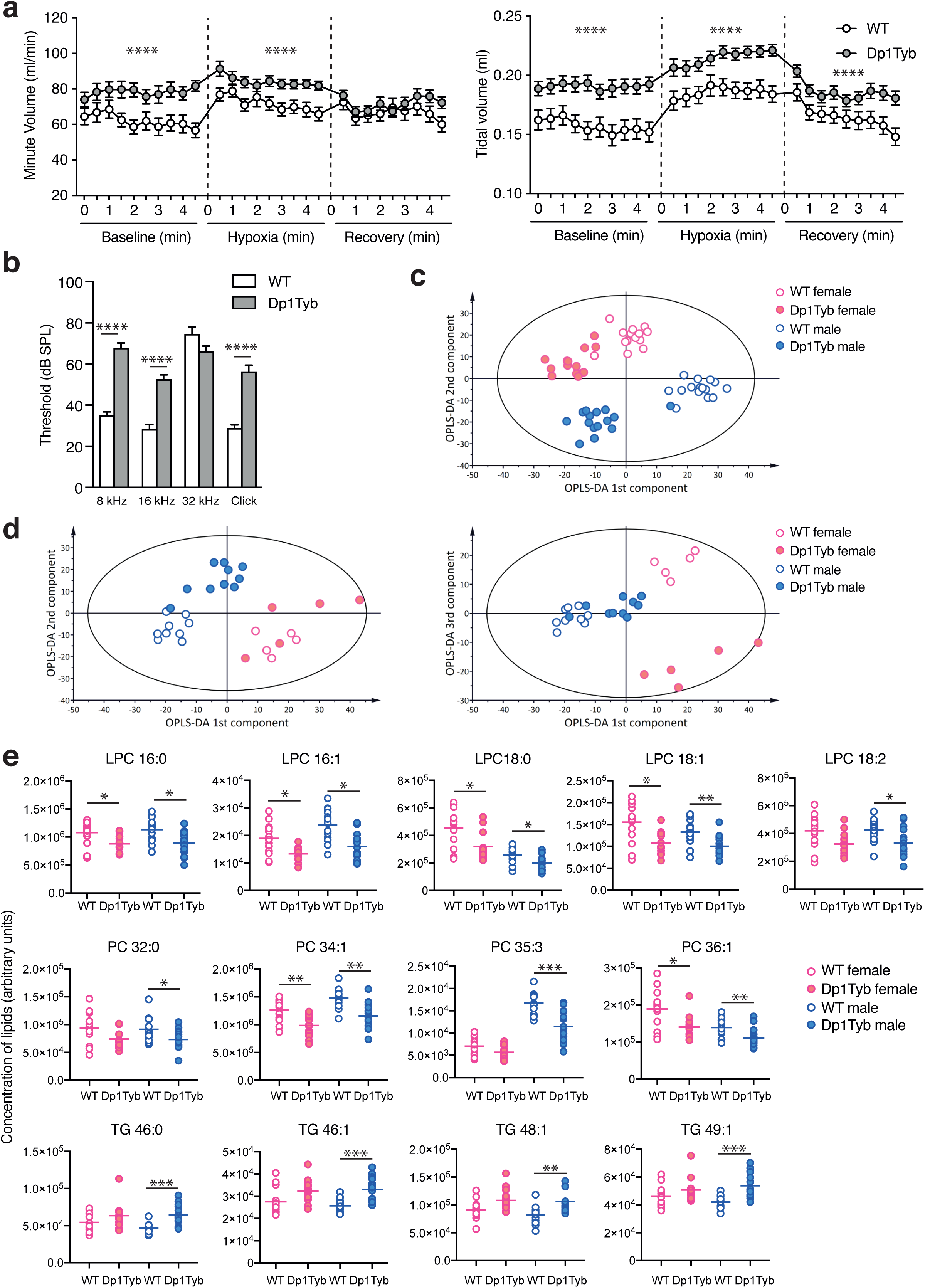
Altered breathing, auditory brainstem response and plasma lipids in Dp1Tyb mice. **a**, Mean±SEM volume of breaths over a minute or single breaths (tidal) taken by WT or Dp1Tyb mice (cohort 1, females and males combined) analysed by whole body plethysmography over three periods of 4 min: baseline normoxia, hypoxia challenge (10% O_2_, 3% CO_2_) and recovery in normoxia. **b**, Mean±SEM auditory brainstem response of WT or Dp1Tyb mice from cohort 1 (females and males combined) at different frequencies (8 kHz, 16 kHz and 32 kHz) and to a click box of mixed tones. **c**, **d**, Graphs of orthogonal partial least squares discriminant analysis (OPLS-DA) of lipids in plasma from (c) fasted (cohort 2) and (d) free-fed mice (cohort mice showing plots of 1^st^ v 2^nd^ or 1^st^ v 3^rd^ OPLS-DA components. The Hotelling T^2^ ellipse indicates the area within which 95% of the samples are expected to lie. **e**, Plasma levels of the indicated lipids (arbitrary units) in fasted WT and Dp1Tyb mice (cohort 2). Horizontal lines indicate mean. * 0.01 < *q* < 0.05; ** 0.001 < *q* < 0.01; *** 0.0001 < *q* < 0.001; **** *q* < 0.0001. LPC, lysophosphatidylcholine; PC, phosphatidylcholine.

### Dp1Tyb mice have impaired hearing

Children with DS often have impaired hearing caused by otitis media, an inflammation of the middle ear^33–35^. To examine if this phenotype was present in Dp1Tyb mice, we evaluated hearing using the auditory brainstem response. We found that compared to WT mice, Dp1Tyb mice had substantially higher minimum sound intensity thresholds required to elicit a brainstem response when challenged with sounds at 8 kHz and 16 kHz and with clicks consisting of mixed frequencies (Fig. 5b). Furthermore, pathological analysis showed that Dp1Tyb mice had otitis media (Supplementary Table 4), which may be causing the impaired hearing.

Children and adults with DS have an increased prevalence of eyesight defects in multiple eye structures, including the lid, iris, cornea, lens and retina^36^. We examined the eyes of Dp1Tyb mice but found no visible defects within the eye (Supplementary Table 5).

### Dp1Tyb mice show characteristics of a pre-diabetic state

DS results in increased prevalence of type 1 and type 2 diabetes^37–40^. Thus, we measured the ability of the mice to clear an intra-peritoneal injection of glucose. This showed no difference between Dp1Tyb and WT mice, thus Dp1Tyb are not diabetic at 13 weeks of age (Supplementary Fig. 3c).

Next, we collected plasma from free-fed and fasted mice, and measured levels of multiple analytes and hormones. We found reduced levels of glucose in fasted but not free-fed Dp1Tyb male mice (Supplementary Fig. 3c, d). Most other analytes were unaffected, except for raised levels of inorganic phosphorus and aspartate aminotransferase (AST), decreased alpha-amylase in free-fed mice, and increased glucagon and adiponectin in fasted mice (Supplementary Fig. 3d, f). The slightly elevated AST/alanine aminotransferase (ALT) ratio (1.66 v 1.19 in Dp1Tyb v WT mice, **Supplementary Table 1**) may indicate liver pathology^41^, but this will require further investigation. Increased adiponectin has been reported in people with DS^42^.

Finally, we used mass spectrometry to measure levels of lipids in plasma from free-fed and fasted mice. Orthogonal partial least squares discriminant analysis (OPLS-DA) of the resulting data showed that for both free-fed and fasted mice, lipid composition was distinct between Dp1Tyb and WT mice and between male and female mice (Fig. 5c, d). The differences in lipid composition were larger for fasted mice, with 85 out of 285 lipids showing a significant difference (q≤0.05), compared to 11 out of 285 lipids in free-fed mice (**Supplementary Table 1**). Dp1Tyb mice had significantly reduced levels of many species of both saturated and unsaturated lysophosphatidylcholine (LPC) and phosphatidylcholine (PC) lipids, and increased levels of many triglycerides (Fig. 5e). Decreased levels of LPC phospholipids and increased triglycerides are associated with a pre-diabetic state^43–47^. Furthermore, higher plasma triglycerides were found in people with non-alcoholic fatty liver disease (NAFLD) and in hepatic steatosis, a clinical subtype of NAFLD^48, 49^. These results suggest that Dp1Tyb mice have characteristics of a pre-diabetic state with associated liver pathology.

### Increased erythropoiesis and megakaryopoiesis in Dp1Tyb mice

Around 10-15% of neonates with DS present with a pre-leukaemic condition known as transient abnormal myelopoiesis (TAM) characterized by an accumulation of megakaryoblasts in the circulation^50, 51^. Most children with TAM undergo spontaneous regression, but in 10-20% of cases, the condition progresses to an acute megakaryoblastic leukaemia known as myeloid leukaemia of DS (ML-DS). Both TAM and ML-DS cells inevitably contain a stereotypical acquired mutation in *GATA1*^50, 51^. While trisomy 21 and *GATA1* mutation are sufficient for the generation of TAM, further mutations are required for progression to ML-DS, typically in cohesin, CTCF or epigenetic regulators such as EZH2. In addition, children with DS have a 20-fold higher risk of developing acute lymphoblastic leukaemia (DS-ALL), usually derived from B-cell progenitors^52^.

Haematological analysis showed that Dp1Tyb mice have reduced numbers of erythrocytes in the blood, a reduced haematocrit, and reduced haemoglobin concentration, but the erythrocytes are larger with more haemoglobin per cell (Fig. 6a). Thus, Dp1Tyb mice have a macrocytic anaemia. Analysis of other haematological parameters showed no changes, except for an increase in the percentage and concentration of monocytes (Fig. 6a, Supplementary Fig. 4).

**Figure 6.**
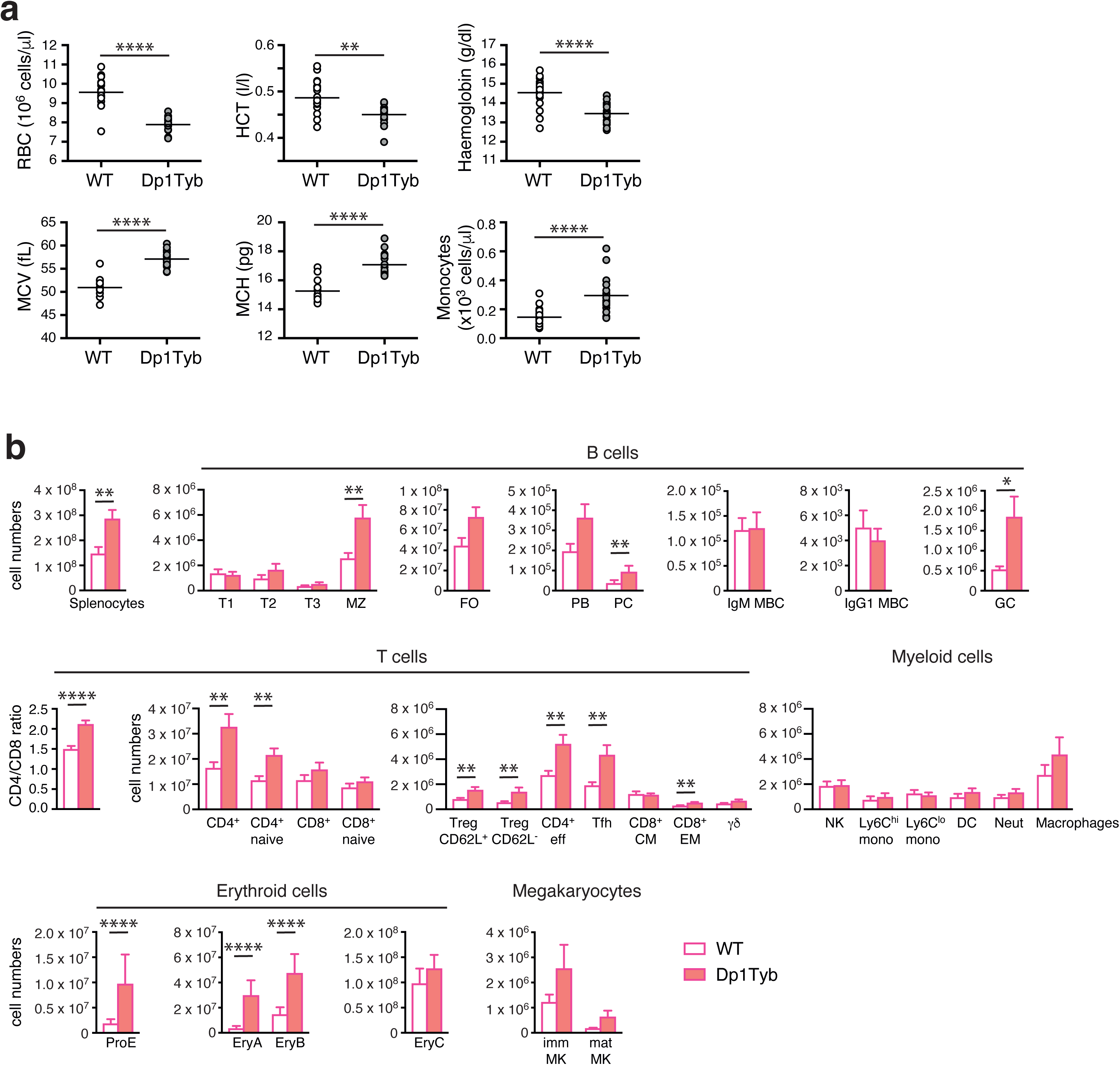
Macrocytic anaemia, splenomegaly, and increased erythropoiesis and megakaryopoiesis in Dp1Tyb mice. **a**, Haematocrit (HCT), mean corpuscular haemoglobin (MCH), mean corpuscular volume (MCV), haemoglobin, monocyte and red blood cell (RBC) concentrations in the blood of WT or Dp1Tyb mice (cohort 1, females and males combined). **b**, Flow cytometric analysis of splenocytes from female WT and Dp1Tyb mice (cohort 4), showing mean±SEM numbers of splenocytes, transitional type 1 (T1), T2, T3, marginal zone (MZ), follicular (FO), germinal centre (GC) B cells, plasmablasts (PB), plasma cells (PC), IgM and IgG1 memory B cells (MBC), mean±SEM ratio of CD4^+^/CD8^+^ T cells, and mean±SEM numbers of total or naive CD4^+^ or CD8^+^ T cells, CD62L^+^ or CD62L^-^ regulatory T cells (Treg), effector T cells, T follicular helper (Tfh) cells, CD8^+^ central memory (CM) and effector memory (EM) T cells, γδ T cells, NK cells, Ly6C^hi^ and Ly6C^lo^ monocytes (mono), dendritic cells (DC), neutrophils (Neut), macrophages, pro-erythroblasts (ProE), EryA, EryB and EryC erythroid progenitors, and immature (imm) and mature (mat) megakaryoblasts (MK). * 0.01 < *q* < 0.05; ** 0.001 < *q* < 0.01; *** 0.0001 < *q* < 0.001; **** *q* < 0.0001.

We next carried out comprehensive flow cytometric analysis of haematopoietic cells in the bone marrow, spleen, lymph nodes, peritoneal cavity and blood of Dp1Tyb and control mice (Supplementary Fig. 5, 6). Bone marrow of Dp1Tyb mice had unchanged numbers of erythroid progenitors and immature and mature megakaryocytes (Supplementary Fig. 7a), but reduced percentages of developing B-lineage subsets (pro-B, pre-B, immature and mature B cells) (Supplementary Fig. 7b). Dp1Tyb mice had an increased number of splenocytes (Fig. 6b), in keeping with the increased splenic size (Supplementary Fig. 1d). Many lymphoid subsets were altered in spleen, including increased numbers of marginal zone B cells, germinal centre B cells, plasma cells, and multiple subsets of CD4^+^ T cells – naïve, effector, regulatory and T follicular helper T cells (Fig. 6b).However, the largest changes in the spleen were substantial increases in the numbers and percentages of pro-erythroblasts (ProE), EryA and EryB erythroid progenitors and immature and mature megakaryocytes (Fig. 6b, Supplementary Fig. 8), explaining the increased extramedullary haematopoiesis of Dp1Tyb spleens (Supplementary Fig. 2a, Supplementary Table 4).

Analysis of blood showed no changes in percentages of leukocytes, including monocytes (Supplementary Fig. 9a), whereas in peripheral and mesenteric lymph nodes of Dp1Tyb mice there were small increases in B cells, and small reductions in CD8^+^ and γδ T cells (Supplementary Fig. 9b, c). In the peritoneal cavity conventional B2 cells were increased at the expense of reduced percentages of B1a cells (Supplementary Fig. 9d). Finally, analysis of developing T cell subsets in the thymus showed that Dp1Tyb mice had reduced numbers and percentages of the early double negative 1 (DN1), DN2 and DN4 cells, but increased numbers of intermediate and mature CD8^+^ single positive thymocytes (Supplementary Fig. 10a, b). Thus, Dp1Tyb mice have increased erythropoiesis and megakaryopoiesis in the spleen and increases in some B and T cell subsets. Other lymphoid tissues showed small changes in B and T cell subsets.

### Dp1Tyb mice have impaired short-term associative memory and disrupted sleep

DS is the most common genetic cause of intellectual disability^53^, arising from cognitive impairment, with delayed acquisition of many developmental milestones^54^. Children and adults with DS have a decreased ability to learn and show deficits in both short-term and long-term memory. DS results in poorer processing of verbal information, slower language acquisition, impaired attention, poor response inhibition and slower planning of tasks. Notable co-morbidities include higher rates of seizures and sleep disruption characterised by poor sleep initiation and maintenance, both of which may contribute to the cognitive impairment.

In the phenotyping pipeline, Dp1Tyb mice were analysed in several behavioural tests. The open field test assesses anxiety, activity and exploratory behaviour. In this test mice are placed in a well-lit arena for 30 min and their activity is monitored, measuring the time they spend in the centre of the arena compared to the periphery, as well as speed of movement and distance travelled. Overall, compared to controls, Dp1Tyb mice visited the centre of the arena less frequently, spent less time in the centre, moved at slower speeds and travelled shorter distances (Fig. 7a). Thus, Dp1Tyb mice are less active and the reduced time spent in the centre indicates that they may be more anxious, although this may also be the result of reduced movement. This analysis was extended using an elevated zero maze which assesses the conflict between exploratory behaviour of novel environments and avoidance of well-lit open areas. This showed no difference between Dp1Tyb and WT mice in the time spent in open versus closed arms of the maze, implying that at least in this assay Dp1Tyb mice do not have increased anxiety (Supplementary Fig. 11a).

**Figure 7.**
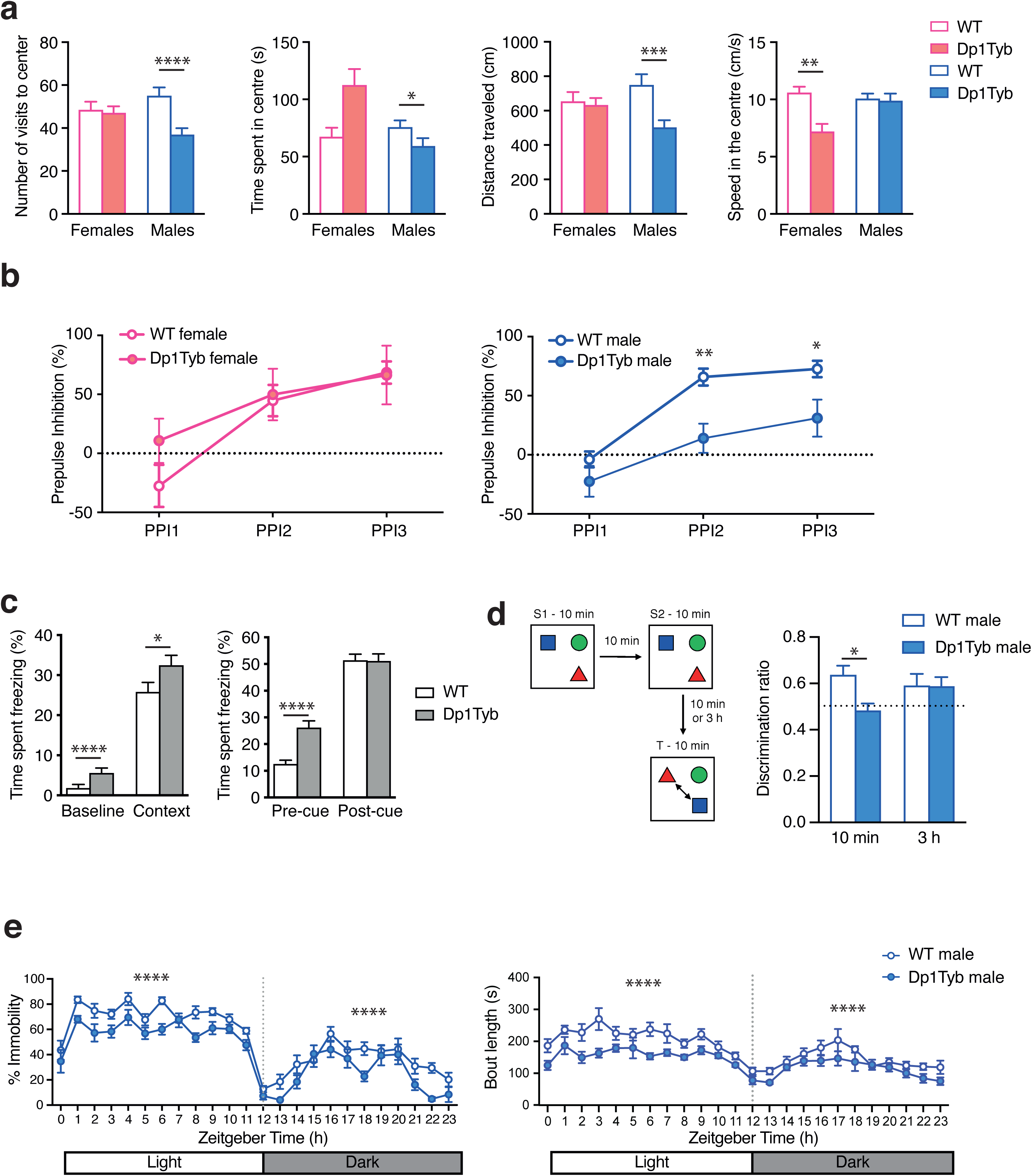
Decreased open field activity, sleep and object-in-place memory in Dp1Tyb mice. **a**, Mean±SEM number of visits in the centre, time spent in the centre, distance travelled in the centre and speed in the centre of mice in cohort 2 undergoing open field habituation on days 1 and 2 of the test. **b**, Mean±SEM pre-pulse inhibition in WT and Dp1Tyb mice (cohort 1) of a response to a startle tone (110dB) to increasing volumes of pre-pulse tones (55, 65, 70 dB; PPI1 to PPI3). **c**, Fear conditioning test on WT and Dp1Tyb mice in cohort 2, showing mean±SEM time spent freezing in response to being placed in chamber associated with aversive experience (context) compared to response to the same chamber before conditioning (baseline), and also freezing time in a novel chamber before or after being given an auditory tone associated with an aversive experience (pre- and post-cue). **d**, WT and Dp1Tyb mice (cohort 6) were placed in an arena with 3 different objects for two 10 min sampling periods (S1, S2), then tested after a delay of either 10 min or 3 h for their ability to recognise which objects had changed location (T, test phase). The discrimination ratio (mean±SEM) measures the preferential interaction with switched objects compared to the unchanged object for mice tested after a 10 min or 3 h delay. A ratio of 0.5 (dotted line) indicates performance at chance level, with no associative object-in-place memory. **e**, Mean±SEM % immobility or length of immobile bouts of WT and Dp1Tyb mice (cohort as a function of time showing the light and dark phases of a 24 h period. Immobility is taken as a proxy of sleep. * 0.01 < *q* < 0.05; ** 0.001 < *q* < 0.01; *** 0.0001 < *q* < 0.001; **** *q* < 0.0001.

Dp1Tyb and control mice were assessed using the pre-pulse inhibition of acoustic startle test, which measures sensorimotor gating mechanisms through exposing the mice to a loud startle sound, which may or may not be preceded by a quieter pre-pulse tone. As the intensity of the pre-pulse tone increases, the neural response to the following startle noise is hindered, resulting in decreased flinching. Such pre-pulse inhibition (PPI) is often impaired in neurological conditions such as schizophrenia and obsessive-compulsive disorder^55^. Both Dp1Tyb and WT mice showed increased PPI with increasing pre-pulse tone intensity, but male Dp1Tyb mice were significantly impaired in this response (Fig. 7b). Since Dp1Tyb mice have impaired hearing (Fig. 5b), it is unclear if the decreased PPI is due to this sensory deficit or to defective sensorimotor gating.

We investigated distinct learning and memory modalities using a number of standard tests. In a fear conditioning test, we assessed memory of an aversive foot shock experience, determining if the animal can associate it with context, a novel chamber, or with a cue in the form of an auditory conditioned stimulus. Responses to the context or cue were measured by the time the animal spent freezing. We found that Dp1Tyb mice responded at least as well as WT mice to both context and cue, indicating that they had no defect in associative memory of the aversive response (Fig. 7c). However, Dp1Tyb mice spent more time freezing when placed into novel chambers (baseline and pre-cue responses) again suggesting that they may be more anxious.

We measured spatial memory using an object-in-place test. The mice were first habituated to a novel arena and then presented with three distinct objects during a sample phase for 10 min, removed for 10 min, and placed back in the arena with the three objects for a further 10 min. Importantly, there were no differences between groups in object contact times during the sample phase. The mice were then removed for either 10 min or 3 h before being placed back in the arena for the test phase in which the locations of two of the objects had been switched. Contact times with objects that had been moved were compared to contact time with the object that had not been moved using a discrimination ratio. We found that Dp1Tyb mice performed worse than WT mice when placed back into the arena after a 10 min delay – unlike WT mice, their discrimination ratio was not significantly different from chance (0.5) (Fig. 7d). In contrast, Dp1Tyb mice performed as well as WT mice after a 3 h delay. Thus, in this paradigm, Dp1Tyb mice have impaired short-term (10 min) but not longer-term (3 h) associative recognition memory.

People with DS often experience sleep disturbance^53^. We assessed sleep in Dp1Tyb mice by tracking movement over a 24 h period in a 12 h light/12 h dark cycle, scoring cumulative periods of immobility of >40 s, which show very high correlation with sleep^56^. Compared to WT mice, this cumulative immobility score was substantially reduced in Dp1Tyb, while immobility bout lengths were significantly shorter, particularly during the light phase where most sleep is scored (Fig. 7e). Thus, Dp1Tyb mice sleep less than WT controls and show a higher number of short-duration bouts, indicating a disrupted sleep pattern.

### Dp1Tyb mice have impaired motor function

Children with DS have impaired motor skills and postural control^57–59^. We assessed multiple aspects of motor function in Dp1Tyb mice using diverse assays. Dp1Tyb mice spent less time in spontaneous wheel running, carrying out fewer runs and rotations, running at lower speeds and for shorter distances (Fig. 8a). In the Locotronic test where mice walk along a horizontal ladder with evenly spaced rungs, Dp1Tyb mice made a larger numbers of errors, indicating defects in motor coordination (Fig. 8b). Defective movement of Dp1Tyb mice was also seen in the CSD test (Fig. 8c). Dp1Tyb mice had normal grip strength indicating that altered locomotor activity was not due to altered muscle tone (Supplementary Fig. 11b). Thus, Dp1Tyb mice have substantially impaired motor function.

**Figure 8.**
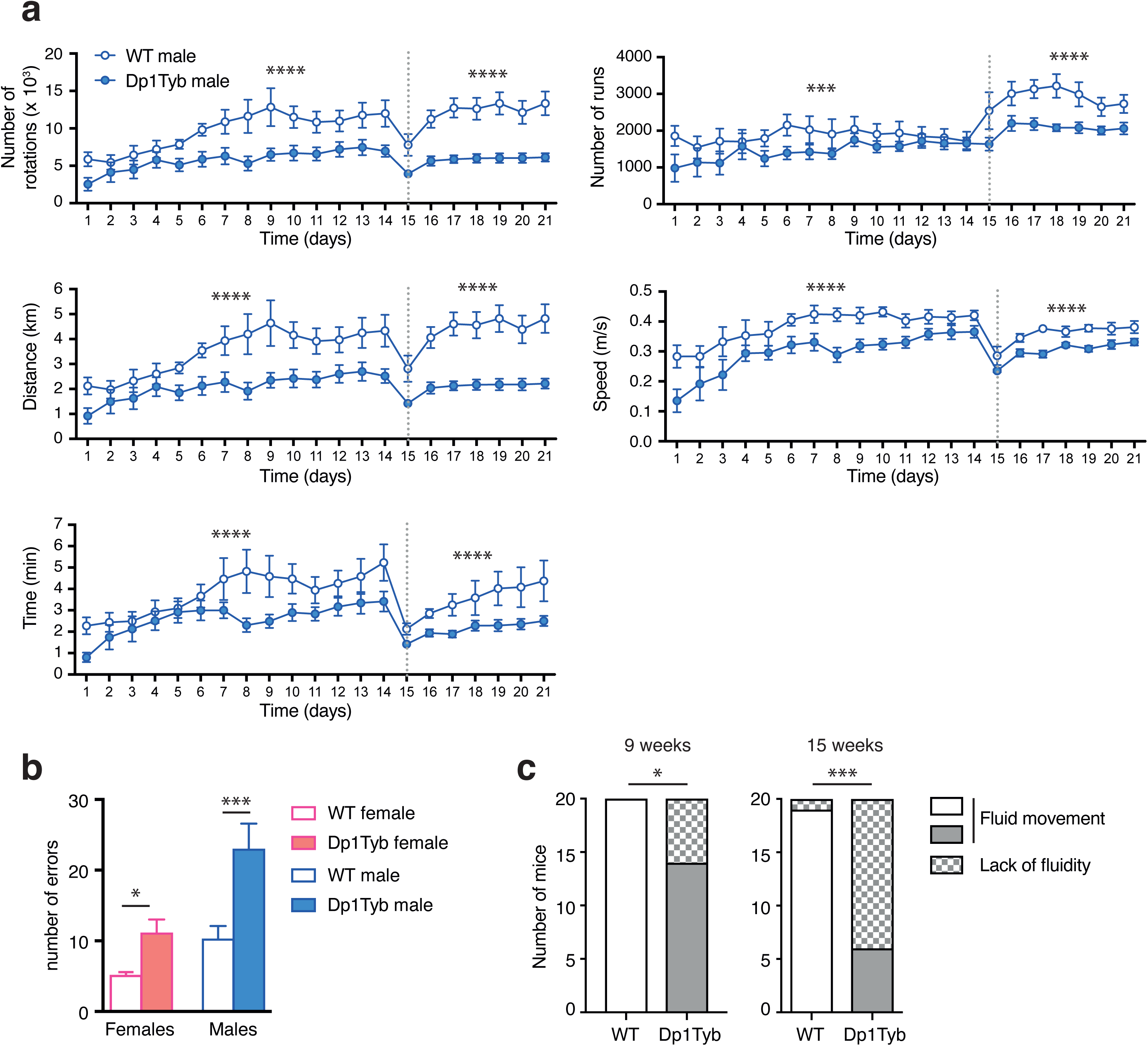
Impaired locomotor activity in Dp1Tyb mice. **a**, Analysis of home cage wheel running by WT and Dp1Tyb mice (cohort 2) over a 3-week period, showing mean±SEM number of rotations, number of runs, distance run, speed and time spent running. Dotted line indicates transition from a simple to a complex wheel in which several rungs were missing. **b**, Mean±SEM number of foot errors made by WT and Dp1Tyb mice (cohort 3) traversing a horizontal ladder in a Locotronic apparatus. **c**, Graph of numbers of WT and Dp1Tyb mice (cohort 1, females and males combined) showing or lacking fluid movement as determined during a SHIRPA test at 9 and 15 weeks of age. * 0.01 < *q* < 0.05; *** 0.0001 < *q* < 0.001; **** *q* < 0.0001.

## Discussion

Analysis of Dp1Tyb mice shows that they had significant differences compared to WT mice in 468 out of 1800 parameters across 22 different tests. Thus, an extra copy of 148 genes has substantial phenotypic consequences in many physiological domains. The reduced perinatal viability of Dp1Tyb pups may be caused by congenital heart defects. At E14.5, around 25% of Dp1Tyb embryos have AVSD, and a further 35% have VSD^11^, similar to the defects reported in around 40% of babies with DS. It is likely that pups with AVSD will die perinatally potentially accounting for the reduced viability.

Many of the phenotypes in Dp1Tyb mice resemble those reported in people with DS. Analysis of skeletal structures, showed that Dp1Tyb mice have reduced bone density, shorter tibia and altered craniofacial structures, in particular brachycephaly. All of these changes are seen in people with DS^23–25, 28, 60^. In agreement with this, Dp1Tyb mice have been reported to have shorter body and femur length^61^ and similar craniofacial changes have been seen in the Dp1Yey mouse strain, which has an extra copy of the same genes as Dp1Tyb^62, 63^

ECG studies in people with DS without congenital heart defects showed increased T-wave length, similar to that seen in Dp1Tyb mice^64^. Cardiac function was altered in 12-week old Dp1Tyb mice with a slower heart rate, and increased cardiac output, stroke volume and left ventricular diameters. Increased cardiac output and stroke volume could be caused by abnormal fluid loading from a left-right shunt such as a septal defect. However, this would be expected to cause an increased heart rate. A more likely explanation is that a shunt during embryonic development would have reduced the effective stroke volume leading to renal hypoperfusion and compensatory salt and water retention to increase circulatory volume. If the shunt closed perinatally, the increased circulatory volume would lead to the observed increased stroke volume and cardiac output with a reactive lower heart rate. Subsequent diuresis may have normalised the circulatory volume thereby resulting in normal cardiac function in the older 57-week old mice.

Lipidomic and clinical chemistry analysis of plasma showed that Dp1Tyb mice have characteristics of a pre-diabetic state with hepatic steatosis, conditions that are seen in people with DS^65^. Further work is needed to determine if the animals could be induced to develop glucose intolerance, for example by being fed a high-fat diet.

Dp1Tyb mice have a macrocytic anaemia alongside increased splenic erythropoiesis, and megakaryopoiesis, resembling a pre-leukaemic condition. However, the animals do not develop overt megakaryoblastic leukaemia, most likely because progression to leukaemia requires mutations in *Gata1* and other genes. Similar macrocytic anaemia and elevated megakaryopoiesis have been reported in several other mouse DS models carrying an extra copy of different sets of Hsa21-orthologous Mmu16 genes^66–, 68^. We detected no increase in B-cell progenitors in Dp1Tyb mice and no DS-ALL, however this leukaemia also requires additional mutations^52^.

Sensory deficits are often seen in DS, including in hearing and vision^53^. Dp1Tyb mice have defective hearing, which is most likely due to otitis media, a common condition in DS^33–35^. We found no eye defects in the mice, but further studies would be needed to evaluate visual acuity. In a previous study we found no defects in nociception or proprioception in Dp1Tyb mice^69^.

Dp1Tyb mice have defective short-term memory, disrupted sleep and motor deficits, neurological features that have also been reported in people with DS^53, 57–59^. Previously we showed that these animals have reduced theta wave frequency in the medial pre-frontal cortex and the hippocampus, and increased theta to high gamma phase amplitude coupling in the hippocampus, along with deficits in spatial working memory^70^. Memory and motor deficits have also been reported in the related Dp1Yey mice^69, 71–73^; the motor deficits in these mice may be partly due to motor neuron loss, which we also found in human DS^69^. The disrupted sleep of Dp1Tyb mice may have a neurological basis, but analysis of Dp1Yey mice showed reduced upper airway volume, suggesting that apnoea may affect sleep in these animals^63^.

While Dp1Tyb mice have many DS-like phenotypes, some are not seen in these animals. AD in DS is most likely caused by a third copy of the *APP* gene on Hsa21^2^. Dp1Tyb mice have three copies of *App* but show no deposition of Aβ in the hippocampus, thus they do not model AD pathology. This is not surprising, because amyloid pathology is only seen in mice expressing mutant amyloidogenic human APP proteins; overexpression of wild-type mouse APP is not sufficient^74^.

Muscle hypotonia and increased body fat are features of DS^27, 75^, but neither was seen in Dp1Tyb mice. Furthermore, Dp1Tyb mice have higher VO_2_, VCO_2_, breathing rates and a higher RER, in contrast to people with DS who show lower VO_2_ and VCO_2_, and reduced breathing rates and RER^76^. Thus, Dp1Tyb mice do not model the reduced cardiorespiratory function in DS. These ‘missing’ phenotypes may be due to increased dosage of Hsa21 genes whose orthologues are in the Mmu10 and Mmu17 regions. To investigate this, a broad phenotypic analysis will be needed of mice that have extra copies of these regions^20^.

The sexually dimorphic nature of some phenotypes should be noted for therapeutic strategies. We discovered several pre-disease states in these mice at 17 weeks of age; it will be important to extend this phenotypic analysis across the lifetime trajectory of the animals, comparing it to human DS. Finally, we note that this wide range of phenotypes was detected on an inbred C57BL/6J background. Since different genetic backgrounds can modify phenotypes, it will be important to broaden the work to other backgrounds.

In summary, we have performed the first comprehensive phenotypic analysis of a mouse model for DS, revealing that Dp1Tyb mice have a plethora of DS-like phenotypes and thus can be used to investigate complex underlying pathological mechanisms. Importantly, the existence of a complete panel of mouse strains with extra copies of shorter regions of Mmu16 will allow mapping and identification of the causative dosage-sensitive genes^11^.

## Acknowledgments

We thank the Flow Cytometry and Biological Research Facility of the Francis Crick Institute for flow cytometry and animal husbandry respectively and Probir Chakravarty for help with bioinformatics analysis. We thank Timothy Dawes for useful discussions. VLJT and EMCF were supported by the Wellcome Trust (grants 080174, 098327 and 098328) and VLJT was supported by the UK Medical Research Council (Programme U117527252) and by the Francis Crick Institute which receives its core funding from Cancer Research UK (FC001194), the UK Medical Research Council (FC001194), and the Wellcome Trust (FC001194). HC, PK-B and SW were supported by the EC FP7 Infrafrontier-13 project (grant number 312325). SG, MS and AMM were supported by the Medical Research Council (MC_U142684171) and National Human Genome Research Institute of the National Institutes of health (UM1HG006370). This research was funded in part by the Wellcome Trust (grants 080174, 098327, 098328 and FC001194). For the purpose of Open Access, the author has applied a CC-BY public copyright licence to any Author Accepted Manuscript version arising from this submission.

## Author contributions

EL-E, HC, SW-S, JM-W, DG, MN, AS, TH, PK-B, CLS, ER, GTB, HM, TC, JT and SN performed experiments; EL-E, HC, SG, JM-W, TH, GTB, HM, TC, MLS, PMN, JG, MG, MS, and VLJT analysed data; HC, PMN, JG, MG, MS, A-MM, SW, EMCF and VLJT supervised the work; EL-E and VLJT wrote the manuscript.

## METHODS

### Mice

C57BL/6J.129P2-Dp(16Lipi-Zbtb21)1TybEmcf (Dp1Tyb) mice^11^ were bred in specific-pathogen free conditions either at the Mary Lyon Centre (MRC Harwell) or at the Francis Crick Institute by crossing mutant mice to C57BL/6J mice. All mice analysed had been backcrossed to the C57BL/6J background for at least 10 generations. Mice in cohorts 1 and 2 were weighed once a week for the duration of each phenotyping pipeline. All tests were carried out by experimenters who were blind to the genotype of the mouse. Zbtb20^Tg^(PDGFB–APPSwInd)^20Lms^ mice (J20)^77^ were bred on a C57BL/6J background at the UCL Institute of Neurology. All animal work was carried out under Project Licences granted by the UK Home Office.

### Analysis of RNA sequencing data

Using our RNA sequencing data from the hippocampi of 5 WT and 5 Dp1Tyb mice aged 18.5-19 weeks^78^, we calculated the fold change in expression for the genes in the duplicated region on Mmu16 (*Lipi* to *Zbtb21*), using the TPM (transcripts per million reads) measure, excluding genes whose expression summed over all 10 samples was < 1 TPM. Significantly differentially expressed genes were identified using DEseq2^79^.

### International Mouse Phenotyping Consortium (IMPC) pipelines

The pipeline of tests for cohort 1 is closely based on those used by the IMPC pipeline^21^. More detailed descriptions of these protocols are available on the IMPRESS website (www.mousephenotype.org/impress).

### X-ray analysis

X-Ray images of the mice were collected whilst the mice were anaesthetised with isoflurane. A lateral view, a dorsal-ventral view and a skull image were all taken to enable a full qualitative assessment of the integrity of the skeleton. A 2 cm lead bar was utilised to provide a calibration scale for the measurement of the tibia length.

### Combined SHIRPA and dysmorphology analysis (CSD)

The combined SHIRPA and dysmorphology test identifies physical and behavioural abnormalities through observation^29^. Mice were individually observed in a series of environments to test a range of attributes including hearing, visual placement, activity, motor coordination, righting ability as well as morphological features.

### DEXA

The body composition of the mice was assessed using the PIXImus Dual Energy X-Ray Absorption machine (GE Medical Systems, USA). Mice were anaesthetised with ketamine/xylazine and high energy X-Ray images were automatically analysed for fat tissue content, lean tissue content, bone mineral density and bone mineral content. While anaesthetised, the mice also underwent the auditory brain stem response test (see below).

### ECHO-MRI

Body composition of the mice was assessed using an EchoMRI whole body composition analyser (Echo Medical System, Houston, USA). The analysis output quantified fat mass, lean mass and water content of the mice.

### Histopathology

Histopathology was performed on all major organs of four male and four female mice of each genotype (WT and Dp1Tyb) at the end of the cohort 1 pipeline. Following necropsy, heart, spleen, and kidneys were weighed. These and other tissues were fixed in 10% neutral buffered formalin, wax embedded, sectioned and stained with haematoxylin and eosin. Slides were reviewed by two veterinary pathologists. Significant findings were scored using a non-linear semi-quantitative approach ^80^.

### Histology for Aβ

Immediately following euthanasia, the brain was removed and immersion fixed in 10% buffered formal saline (Pioneer Research Chemicals, UK). After 48-72 h, the brain was blocked into 3 mm coronal slices using an adult mouse brain matrix and slices were embedded in paraffin wax using a Sakura VIP6 Automated Vacuum Tissue Processor. A series of 4 μm sections comprising the dorsal hippocampus were cut and mounted onto SuperFrost Plus glass slides. For amyloid-β immunostaining, sections were dewaxed, rehydrated through an alcohol series to water, pre-treated with 80% formic acid for 8 min followed by washing in distilled water for 5 min. Sections were loaded as wet mounts into a Ventana Discovery XT automated stainer, where further pre-treatment for 30 min with mild CC1 (EDTA Boric Acid Buffer, pH 9.0) and blocking for 8 min with Superblock (Medite, #88-4101-00), were performed prior to incubation for 8 h with biotinylated mouse monoclonal antibody 4G8 (2 μg/ml, Sigma-Aldrich SIG-39240 Beta-Amyloid). Staining was completed with the Ventana XT DABMap kit and a haematoxylin counterstain, followed by dehydration and permanent mounting with DPX. All images were acquired using a Leica DM2000 LED microscope fitted with a MC190 HD camera.

### Calorimetry

The metabolic rate of the mice was assessed using indirect calorimetry. Mice were individually housed overnight for 21 h in Phenomaster cages (TSE Systems, Germany) with standard bedding and igloos. Oxygen consumption (VO_2_) and carbon dioxide production (VCO_2_) were simultaneously measured through an indirect gas calorimetry system air sampling, and from this the respiratory exchange ratio (VCO_2_/VO_2_) and heat production were calculated. Activity was monitored using a photobeam-based system from which speed of movement of the animal was calculated.

### Echocardiogram and ECG

The cardiac phenotype of the mice was assessed using both echocardiogram and electrocardiogram (ECG). For mice in cohort 1, these procedures were conducted at the same time. Mice were anaesthetised under isoflurane and the ECG was recorded using BioAmp (AD Instruments, Australia) and LabChart Pro software (AD Instruments). For the echocardiogram, the left ventricle of the heart was imaged and analysed using the Vevo770 (Fujifilm Visualsonics, USA).

### Plethysmography

Mice were individually placed into whole body plethysmography chambers (Data Sciences International) and allowed to acclimatise for 30 min. Breathing volumes of the mice were recorded initially for 5 min in room air (baseline, normoxia), then for 5 min in 10% O_2_/3% CO_2_ (hypoxia) and a further 5 min in room air (recovery, normoxia). Any mice experiencing breathing problems were removed from the chambers immediately.

### Auditory brain stem response

To determine their hearing range, mice were anaesthetised with ketamine/xylazine and the auditory brain stem response was recorded in response to either a click sound or to tones at 8 kHz, 16 kHz and 32 kHz, using subdermal electrodes placed on the vertex and the left and right bulla^81, 82^. Intensity of the sounds was increased from 0 dB to 85dB sound pressure level (SPL) and the threshold of the response was determined as the lowest sound intensity that gave a recognisable ABR waveform response.

### Opthalmoscopy

Eye morphology and visual response of the mice was assessed using a slit lamp and an opthalmascope. Tropicamide was used to dilate the pupils and observations were manually scored for morphological or response abnormalities.

### Glucose tolerance test

Mice were fasted overnight for 18 h. A sample of blood was analysed from the tail, to determine the fasted blood glucose concentration using the Accu-Chek glucose meter (Abbott, UK). The mice were injected intraperitoneally with 20% glucose solution (2 g glucose/kg body weight). Blood glucose measurements were taken again at 15, 30, 60 and 120 min after injection of glucose.

### Blood collection

Mice were anaesthetised with isoflurane and blood collected under anaesthesia from the retro-orbital sinus into either Lithium-Heparin coated tubes for clinical chemistry and into EDTA-coated tubes for haematology. The mice were either free-fed (cohort 1) or fasted overnight for 18 h (cohort 2).

### Clinical chemistry and ELISA

Lithium heparin samples from both free-fed and fasted mice were kept on wet ice and centrifuged within 1 h of collection for 10 min at 5,000 x g in a refrigerated centrifuge set to 8°C. The resulting plasma samples were frozen until analysis. Clinical chemistry of free-fed plasma was analysed with a Beckman Coulter AU680 clinical chemistry analyser using reagents and settings recommended by the manufacturer for the analysis of alanine aminotransferase (ALT), albumin, alkaline phosphatase (ALP), alpha-amylase, aspartate aminotransferase (AST), calcium, chloride, creatinine kinase, free fatty acids, fructosamine, glucose, glycerol, HDL-cholesterol, inorganic phosphorus, iron, potassium, sodium, total bilirubin, total protein, triglycerides and urea. In addition, glucose and triglycerides were also measured in plasma from fasted mice. Insulin, glucagon, leptin and adiponectin were measured in plasma from fasted mice using ELISA kits from Millipore (EZRMI-13K), Mercodia (10-1281-01), Biovendor (RD291001200R) and R&D Systems (MRP300) respectively.

### Lipidomic analysis of plasma

To profile the lipidome, 15 µL plasma from either free-fed or fasted mice was analysed. Lipids were extracted using 250 µL methanol, sonicated, centrifuged and the supernatant was dried under nitrogen. Extracts were reconstituted in 150 µL of 1/2 (v/v) methanol/water and 2 µL was analysed using a Waters Xevo G2 Quadrupole Time of Flight (Q-ToF) mass spectrometer (MS) connected to a Waters Acquity ultra-performance liquid chromatogram (UPLC) (Milford, MA, USA). Chromatography was performed using a Waters Acquity UPLC CSH C18 1.7 µm, 100 x 2.1 mm column, a linear gradient consisting of solvent A (10 mM ammonium formate in 60/40 (v/v) acetonitrile/water) and solvent B (10 mM ammonium formate in 10/90 (v/v) acetonitrile/propan-2-ol) (flow rate = 0.4 mL/min; temp = 55°C; starting conditions 45% B, 7.5 min 98% B). Ions were detected in positive mode with a source temperature of 12°C, desolvation temperature of 550°C, capillary voltage of 1.5 kV, cone voltage of 30 V, cone gas flow of 50 L/h and desolvation gas flow of 900 L/h. Data was converted to netCDF format and the matchedFilter peak-finding algorithm of the xcms software^83^ was used to identify mass peaks. Annotation of lipids was based on exact mass using the LipidMaps database (https://lipidmaps.org/), fragmentation and chromatographic retention time. Data was processed using OPLS-DA within the SIMCA package (Umetrics, Umea, Sweden) and univariate statistics as described below.

### Haematology

Full blood counts and differential analyses of whole blood samples collected from free-fed mice (cohort 1) in EDTA-coated tubes were performed with a Siemens Advia 2120 analyser using reagents and settings recommended by the manufacturer.

### Flow Cytometry

Single cell suspensions from spleen, bone marrow, thymus, mesenteric and peripheral lymph nodes, peritoneal cavity and blood were depleted of erythrocytes using ACK lysis buffer as previously described^84^, before staining cells with a mixture of antibodies in PBS containing the live/dead marker Zombie Aqua (BioLegend). Antibodies against the following antigens were used [indicating antigen-fluorophore (clone)]: B220-BV605 (RA3-6B2), B220-FITC, B220-PE, CD3ε-PE (145-2c11), CD11b-FITC (M1/70), CD11c-PerCP/Cy5.5 (N418), CD19 APC (1D3), CD23-APC (B3B4), CD38-PE (90), CD42d-APC (1C2), CD62L-BV711 (MEL-14), CD49b-FITC (Dx5), CD115-PE (AFS98), CD138-PE/cy7 (281-2), CXCR5-BV785 (L138D7), Gr-1-PE (RB6-8C5), GR-1-FITC, Ly6C-BV711 (HK1.4), Ly6G-FITC (1A8), PD-1-PE/Cy7 (29F.1A12), TCRγδ-BV605 (GL3) and Thy1.2-BV605 (30-H12) from BioLegend; B220-APC/eF780, CD2-PE (RM2-5), CD3ε-FITC, CD4-APC (RM4-5), CD5-PE (53-7.3), CD8-FITC (53-6.7), CD8-PE, CD11b-eF450, CD21-APC/eF780 (4E3), CD25-PE (PC61.5), CD44-APC (IM7), CD71-PE (RI7 217.1.4), CD73-PE/Cy7 (eBioTY-11.8), CD93-APC (AA4.1), Fas-PE/Cy7 (Jo), F4/80-APC (BM8), IgD-eF450 (11-26), NK1.1-PE/Cy7 (PK136) and TCRβ-PerCP/Cy5.5 (H57-597) from eBioscience; CD41-FITC (MWReg30), Fas-PE-CF594, IgG1-APC (X56), PD-L2-BV421 (Ty25) and TER-119-BV421 (TER-119) from BD Biosciences. Goat-anti-mouse IgM Fab-FITC was purchased from Jackson ImmunoResearch. To exclude lineage^+^ cell populations, cells were stained with antibodies against B220, CD3ε, CD11b and Gr-1 for erythrocyte and megakaryoblast staining in spleen and bone marrow, B220 and CD3ε for myeloid cell staining in spleen and blood and B220, CD4, CD8, TCRβ, TCRγδ, Gr-1, CD11b, CD11c, CD49b and NK1.1. for double negative thymocyte staining. An unlabelled antibody against CD16/32 (Fc-block, eBioscience) was used in all stainings to avoid non-specific binding of fluorophore-labelled antibodies to Fcγ receptors. Data was acquired on an LSRII flow cytometer (Becton Dickinson) and analyzed using FlowJo v10.5 (TreeStar).

### Open field habituation

The open field habituation test assesses anxiety, activity and exploratory behaviour in first novel and then familiar environments. Mice were placed in well-lit arenas (150-200 lux, 44 x 44 cm) for 30 min and the activity of the mice in the centre zone (40% of the total area), periphery (area 8 cm towards the centre from the wall) and whole arena was captured in 5 min bins using Ethovision software (version 8.5). Mice were returned to the arena on the following day and measurements repeated to assess habituation.

### Elevated Zero maze

The elevated zero maze assesses the conflict between exploratory behaviour of novel environments and avoidance of well-lit open areas^85^ and uses an elevated maze (53cm off the ground) in the shape of a circle which is divided equally into two open sections and two closed sections, with each section being approximately 30 cm long. Mice were placed onto an open section and allowed to explore for 5 min. The video was analysed using Ethovision software and the total amount of time spent in either the open or closed sections was determined.

### Acoustic startle and pre-pulse inhibition (PPI)

The acoustic pre-pulse startle test measures sensorimotor gating mechanisms through exposing the mice to a loud startle noise which may or may not be preceded by a quieter tone of differing sound levels. As the intensity of the pre-pulse tone increases, the brain should increasingly disregard the following startle noise, resulting in decreased flinching. Mice were placed in an acoustic startle chamber (Med Associates Inc, USA) and acclimatised to a background noise level of 50 dB for 5 min. Mice were then exposed to a startle tone of white noise at 110 dB for 40 ms, either on its own or preceded 50 ms earlier by a 10 ms pre-pulse white noise at 55, 65 or 70 dB (PPI1, PPI2 or PPI3) in pseudorandom order. Responses to the startle tone were measured for 100 ms following the start of the startle tone using a piezoelectric transducer in the floor of the chamber which detected movement of the animal. Each trial condition was tested 10 times.

### Fear conditioning

Fear conditioning assesses the memory of an aversive experience and determines if it is based on the cue, the context or both^85^. Mice were placed into square chambers (17 cm^2^) on day one and allowed to explore for 150 s during which the amount of freezing behaviour was measured (baseline). An auditory stimulus (70 dB) was presented for 30 s and was followed by one foot shock (0.5 mA, 0.5 s). After an interval, the auditory stimulus and foot shock were repeated twice more before the mouse was returned to its home cage. On day 2, the mice were placed into the same chambers and the amount of freezing behaviour was measured over a 5 min period (context). Four hours later, the mice were placed into circular chambers (20 cm in diameter) with additional vanillin essence to reinforce the novel environment setting. Freezing in the new arena was measured for 180 s (pre-cue), after which the auditory stimulus was presented alone and again the amount of freezing behaviour was measured (post-cue). The comparison of baseline v context or pre-cue v post-cue percentage freezing behaviour was used to evaluate associative memory to context or cue respectively.

### Object-in-Place

Mice were first habituated to a transparent Plexiglas arena (60 x 60 x 40 cm) for 10 min for two consecutive days. To test for object-in-place associations, mice were placed in the arena with three distinct objects for 10 min, returned to their home cage for 10 min and brought back into the arena for a further 10 min (sample phases 1 and 2). The mice were returned to their home cage for either 10 min or 3 h and then once again brought into the arena for 10 min, but this time the position of two of the objects in the arena had been exchanged (test phase). Behaviour of mice was recorded with an overhead video camera, and subsequently contact times of the mice with each object were determined; contact was defined by the mouse being ≤ 1 cm from the object and facing towards it. A discrimination ratio was calculated to reflect the preference for contact with the two switched objects compared to the object that had not been moved. The ratio ranged from 0 to 1, ratios over 0.5 indicated a preference for the switched objects, and thus intact object-in-place associative memory.

### Sleep analysis

Analysis of sleep/wake cycles was performed using video tracking as previously described^56^. Briefly, mice were singly housed and placed in light-controlled chambers with near infra-red miniature CCD cameras positioned above the cages (Maplin, UK). Monitoring during dark periods was performed using infra-red illumination. Mice were allowed to acclimatize to the home cage for 24 h in a 12 h light/dark cycle (100 lux light intensity) before data collection. Video monitoring was then performed for a 24 h period in a 12:12 h light/dark cycle. Video files were uploaded to ANY-maze video analysis software (Stoelting, US), which was used to track mouse mobility and to score cumulative periods during which the animals remained immobile for 40 s or more, a measure showing very high correlation with sleep^56^.

### Motor function

Mice were singly housed in cages equipped with modular running wheels (TSE-Systems, Germany)^86^. The average number of rotations, duration of each running bout and distance travelled every night was calculated using TSE-systems software over the course of 21 days. At the end of 14 days, the wheel was changed for a complex wheel in which a number of rungs had been removed, allowing assessment of the ability of the mice to adjust to a more difficult locomotor activity.

### Locotronic test

The Locotronic test (Intellibio, France) is a test of motor co-ordination capability. Mice traverse a horizontal ladder with evenly spaced rungs, along a narrow corridor to reach the exit. The number of rungs that the mouse stepped on or missed was recorded automatically; a missed rung was considered an error. Each animal was tested 2-3 times. Trials where mice took more than 60 s to traverse the ladder were excluded from the analysis; a total of 11 out of 146 trials were excluded, 5 from WT mice and 6 from Dp1Tyb mice, indicating no difference in motivation between genotypes.

### Grip strength

Grip strength was measured using a grip strength meter (Bioseb, France), recording the maximum force generated by a mouse using just its forelimbs or all four limbs. Grip strength measures were carried out in triplicate for each mouse.

### Statistical analysis

For each of the above procedures we took all the phenotype measurements for the Dp1Tyb mice and compared these to values from wild-type control mice. Using fitted linear models the null hypothesis was that a fitted model without a genotype term is as good a fit to the data as a model with the addition of a genotype term. The significance of the good fit is taken as the p-value for a genotype effect between the mutant and wild-types (p(genotype)). In order to adjust for parameters that have non-normal distributions, a Box-Cox procedure^87^ was applied to the data prior to analysis. For continuous variables, a mixed model approach was used to compensate for any batch effects within the data. We constructed a linear-mixed-model of phenotype value as a function of genotype (‘value∼1+sex+experimentrepeat+(1|dop)+genotype’) and compared this with the model without a genotype term (‘value∼1+sex+experimentrepeat+(1|dop)’), where ‘dop’ is the date of procedure. An ANOVA test was used to generate a p(genotype) value for the genotype effect. Where there were no repeated measurements for a parameter, or only a single sex was measured the ‘sex’ and ‘experiment repeat’ formula terms were dropped from the models. For discrete variables an analogous logistical regression model method was used with a model formula similar to the one used for the continuous method but removing the date of procedure mixed effect (as logistical regression does not support it). Once the p(genotype) value was determined for a parameter, the false discovery rate (FDR) adjustment^88^ was used to generate FDR-adjusted q-values: q(genotype). A q-value of ≤0.05 was considered significant, and q(genotype) values are indicated on Figures.

A comparison of variances between the WT and Dp1Tyb data for each parameter was carried out using Brown-Forsythe Levene-type test for equality of variances^89^, which produces a p-value with the null hypothesis that each group has the same variance. FDR adjustment was then carried out on this set of p-values, to produce FDR adjusted q-values for the similarity of variances: q(variance). Sexual dimorphism (sexdim) was assessed by using a mixed-model comparing a model with sex-genotype interaction (‘value∼1+sex+experimentrepeat+(1|dop)+sex:genotype’) with a model without the interaction (‘‘value∼1+sex+experimentrepeat+(1|dop)+genotype’). ANOVA was used to determine the significance of the sexdim and FDR was then applied to this set of p-values to produced q-values, q(sexdim). Metadata fields such as ‘experimenter id’ or ‘anaesthesia’ were analysed to see if the metadata field had a significant effect to the phenotype calls. If the metadata field produced a significant effect on the model (ANOVA, p≤0.05), it was added as a factor to the linear model, compensating for its effect. Coefficient of variation (CoV) for each parameter with continuous variables was calculated by dividing the standard deviation by the mean, treating each sex and genotype separately.

## Supplementary Data

**Supplementary Table 1. Statistical Analysis.**

See Excel file.

**Supplementary Table 2.**
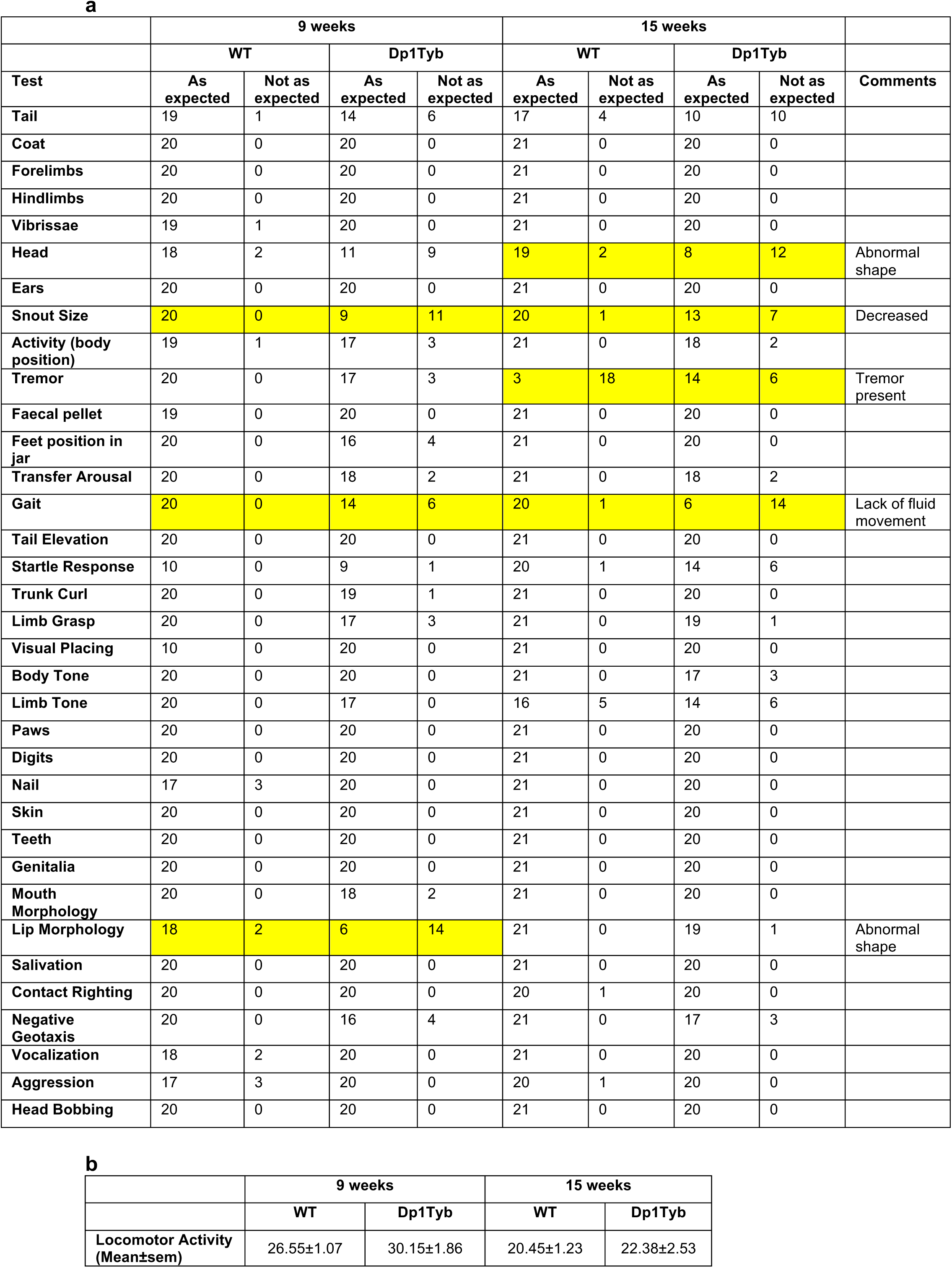
Dp1Tyb mice have abnormal heads, snouts, lips and gait. **a,** Table shows numbers of mice that gave the expected or not expected result in each of the tests in the Combined SHIRPA and Dysmorphology (CSD) tests. Dp1Tyb and WT mice were tested at 9 and 15 weeks of age (cohort 1). Test showing statistically significant differences (q<0.05) between Dp1Tyb and WT mice at either 9 or 15 weeks or both are highlighted in yellow. **b,** Locomotor activity measured as mean±sem number of squares entered by the mouse in 30s.

**Supplementary Table 3.** Dp1Tyb mice have abnormally shaped heads. Table shows numbers of Dp1Tyb and WT mice that were assessed as being normal or abnormal for each of the features listed from X-ray analysis of mice tested at 14 weeks of age (cohort 1). Test showing statistically significant differences (q<0.05) between Dp1Tyb and WT mice is highlighted in yellow.

|  | WT |  | Dp1Tyb |  | Comments |
| --- | --- | --- | --- | --- | --- |
|  | Normal | Abnormal | Normal | Abnormal |  |
| **Skull shape** | 19 | 1 | 0 | 20 | Abnormal shape |
| **Zygomatic bone** | 19 | 1 | 19 | 1 |  |
| **Maxilla** | 19 | 1 | 13 | 7 |  |
| **Clavicle** | 20 | 0 | 20 | 0 |  |
| **Scapulae** | 20 | 0 | 20 | 0 |  |
| **Humerus shape** | 20 | 0 | 20 | 0 |  |
| **Fusion of ribs** | 20 | 0 | 20 | 0 |  |
| **Shape of ribs** | 20 | 0 | 20 | 0 |  |
| **Vertebrae shape** | 19 | 0 | 19 | 0 |  |
| **Fused vertebrae** | 19 | 1 | 19 | 1 |  |
| **Processes on vertebrae** | 20 | 0 | 20 | 0 |  |
| **Joints** | 20 | 0 | 20 | 0 |  |
| **Syndactylism** | 20 | 0 | 20 | 0 |  |
| **Brachydactyly** | 20 | 0 | 20 | 0 |  |
| **Digit integrity** | 20 | 0 | 20 | 0 |  |
| **Pelvis** | 18 | 2 | 16 | 4 |  |
| **Femur shape** | 20 | 0 | 20 | 0 |  |
| **Fibula** | 20 | 0 | 20 | 0 |  |
| **Tibia** | 20 | 0 | 20 | 0 |  |
| **Radius** | 20 | 0 | 20 | 0 |  |
| **Ulna** | 20 | 0 | 20 | 0 |  |
| **Mandibles** | 19 | 1 | 20 | 0 |  |
| **Teeth** | 19 | 1 | 19 | 0 |  |

**Supplementary Table 4.** Dp1Tyb mice have increased splenic extramedullary haematopoiesis, portal tract anomalies in the liver and otitis media. Table shows incidence of tissues with an appearance considered microscopically within normal limits and tissues where pathological findings are present. Pathological findings may be spontaneous background lesions expected in the strain of mouse or considered specific to the genotype of the animals. Pathological analysis on WT and Dp1Tyb mice in cohort 1 was carried out at 16 weeks of age. Significant findings (different from variations in spontaneous background pathology) in Dp1Tyb mice were increased splenic extramedullary haematopoiesis, portal tract anomalies in the liver and otitis media. These are highlighted in yellow.

|  | Females |  |  |  | Males |  |  |  |  |
| --- | --- | --- | --- | --- | --- | --- | --- | --- | --- |
|  | WT |  | Dp1Tyb |  | WT |  | Dp1Tyb |  |  |
| Tissue | Within normal limits | Pathological findings present | Within normal limits | Pathological findings present | Within normal limits | Pathological findings present | Within normal limits | Pathological findings present |  |
| Salivary gland | 4 | 0 | 3 | 1 | 2 | 2 | 5 | 0 |  |
| Lungs | 2 | 2 | 1 | 3 | 4 | 0 | 2 | 3 | Increased perivascular inflammation in Dp1Tyb mice but considered within normal variation in background mouse strain. |
| Thyroid | 4 | 0 | 4 | 0 | 4 | 0 | 5 | 0 |  |
| Trachea | 4 | 0 | 4 | 0 | 4 | 0 | 4 | 1 |  |
| Parathyroid | 2 | 0 | 2 | 0 | 3 | 0 | 2 | 0 |  |
| Kidney | 0 | 4 | 0 | 4 | 2 | 2 | 0 | 5 | Pelvic dilatation and inflammatory cell infiltrates seen in both genotypes. Significance uncertain but considered within normal variation in background mouse strain. |
| Heart | 4 | 0 | 4 | 0 | 4 | 0 | 5 | 0 |  |
| Tongue | 4 | 0 | 4 | 0 | 4 | 0 | 5 | 0 |  |
| Adrenal | 1 | 3 | 1 | 3 | 3 | 1 | 2 | 3 | Subcapsular cell hyperplasia in both genotypes but considered within normal variation in background mouse strain. |
| Liver | 0 | 4 | 0 | 7 | 3 | 1 | 0 | 4 | Significant lesion. Bile duct hyperplasia and prominent portal vasculature increased in Dp1Tyb mice. Not seen spontaneously. |
| Pancreas | 4 | 0 | 5 | 2 | 4 | 0 | 3 | 1 | Fatty atrophy and inflammatory cell foci in Dp1Tyb mice but considered within normal variation in background mouse strain. |
| Gall bladder | 3 | 0 | 6 | 0 | 3 | 1 | 2 | 2 |  |
| Spleen | 0 | 4 | 0 | 7 | 0 | 4 | 0 | 4 | Significant lesion. Extramedullary haematopoiesis increased in Dp1Tyb mice. |
| Thymus | 3 | 1 | 5 | 2 | 4 | 0 | 3 | 1 |  |
| Mesenteric lymph node | 3 | 0 | 2 | 0 | 3 | 0 | 3 | 0 |  |
| Oesophagus | 3 | 1 | 4 | 0 | 4 | 0 | 4 | 0 |  |
| Duodenum | 3 | 0 | 3 | 0 | 4 | 0 | 4 | 0 |  |
| Jejunum | 4 | 0 | 4 | 0 | 4 | 0 | 4 | 0 |  |
| Ileum | 4 | 0 | 4 | 0 | 4 | 0 | 4 | 0 |  |
| Colon | 3 | 1 | 4 | 0 | 4 | 0 | 4 | 0 |  |
| Caecum | 4 | 0 | 4 | 0 | 4 | 0 | 4 | 0 |  |
| Stomach | 4 | 0 | 4 | 0 | 4 | 0 | 4 | 0 |  |
| Bladder | 3 | 0 | 4 | 0 | 4 | 0 | 4 | 0 |  |
| Skin | 4 | 0 | 4 | 0 | 4 | 0 | 4 | 0 |  |
| Muscle | 4 | 0 | 4 | 0 | 4 | 0 | 4 | 0 |  |
| Eye | 4 | 0 | 4 | 0 | 3 | 1 | 4 | 0 |  |
| Brain | 3 | 1 | 2 | 2 | 4 | 0 | 3 | 1 | Ventricular dilatation more frequent in Dp1Tyb mice but considered within normal variation in background mouse strain. |
| Spinal cord | 4 | 0 | 4 | 0 | 4 | 0 | 4 | 0 |  |
| Pituitary | 4 | 0 | 4 | 0 | 3 | 1 | 3 | 1 |  |
| Sciatic nerve | 4 | 0 | 4 | 0 | 4 | 0 | 4 | 0 |  |
| Femur | 4 | 0 | 4 | 0 | 4 | 0 | 4 | 0 |  |
| Sternum | 4 | 0 | 4 | 0 | 4 | 0 | 3 | 0 |  |
| Nasal cavity | 3 | 1 | 3 | 1 | 4 | 0 | 4 | 0 |  |
| Harderian gland | 0 | 0 | 0 | 0 | 4 | 0 | 4 | 0 |  |
| Ear | 0 | 0 | 0 | 2 | 4 | 0 | 0 | 4 | Significant lesion. Otitis media in Dp1Tyb mice. |
| Uterus | 4 | 0 | 4 | 0 |  |  |  |  |  |
| Ovary | 4 | 0 | 4 | 0 |  |  |  |  |  |
| Testes |  |  |  |  | 1 | 3 | 1 | 3 | Tubular atrophy in both genotypes but considered within normal variation in background mouse strain. |
| Seminal vesicles |  |  |  |  | 3 | 1 | 4 | 0 |  |
| Epididymides |  |  |  |  | 4 | 0 | 4 | 0 |  |
| Prostate |  |  |  |  | 4 | 0 | 4 | 0 |  |

**Supplementary Table 5.**
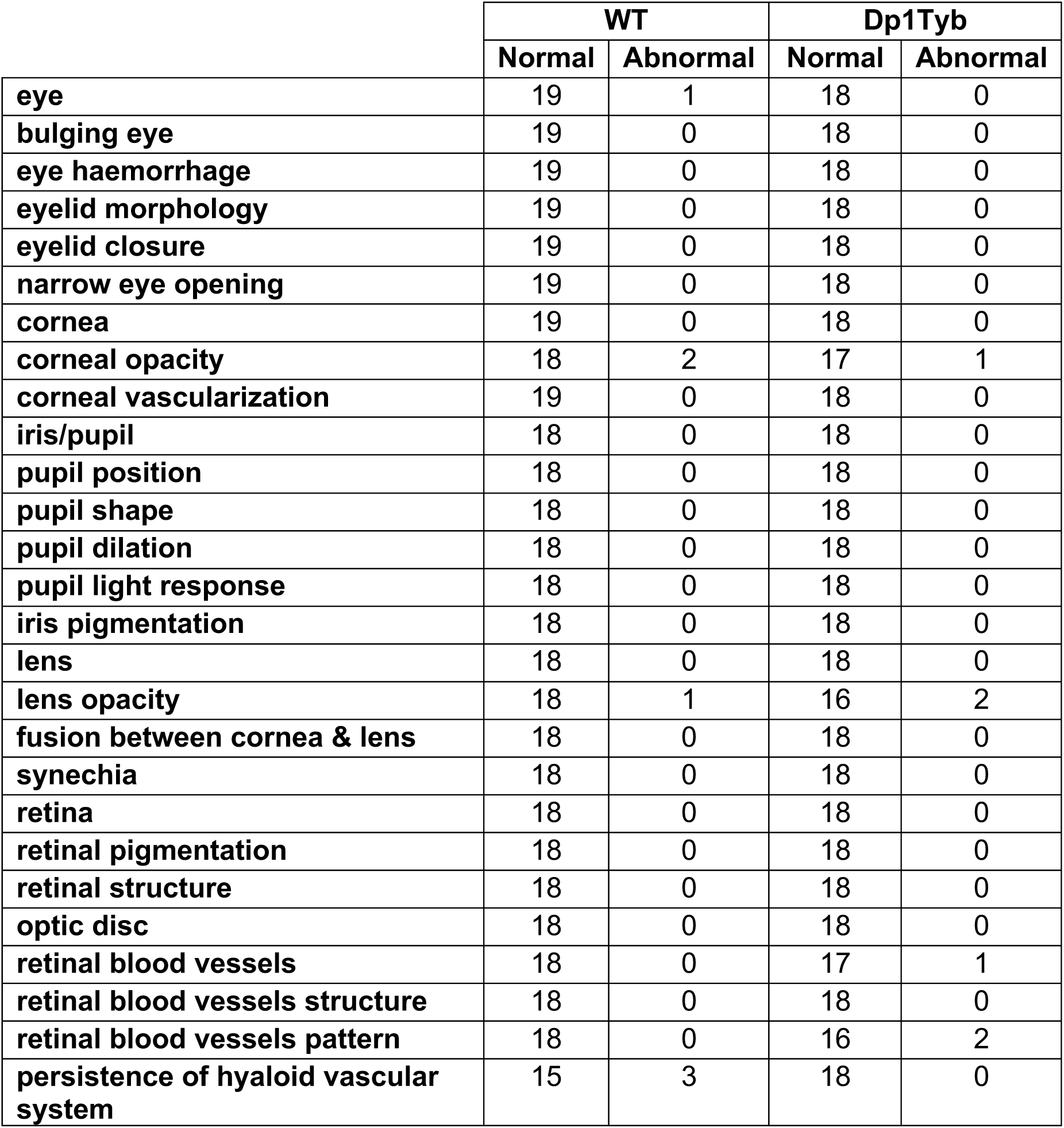
No significant eye abnormalities in Dp1Tyb mice. Table shows numbers of Dp1Tyb and WT mice that were assessed as being normal or abnormal for each of the features or pathologies listed following eye examination at 15 weeks of age (cohort 1). There were no significant differences between Dp1Tyb and WT mice.

**Supplementary Figure 1.**
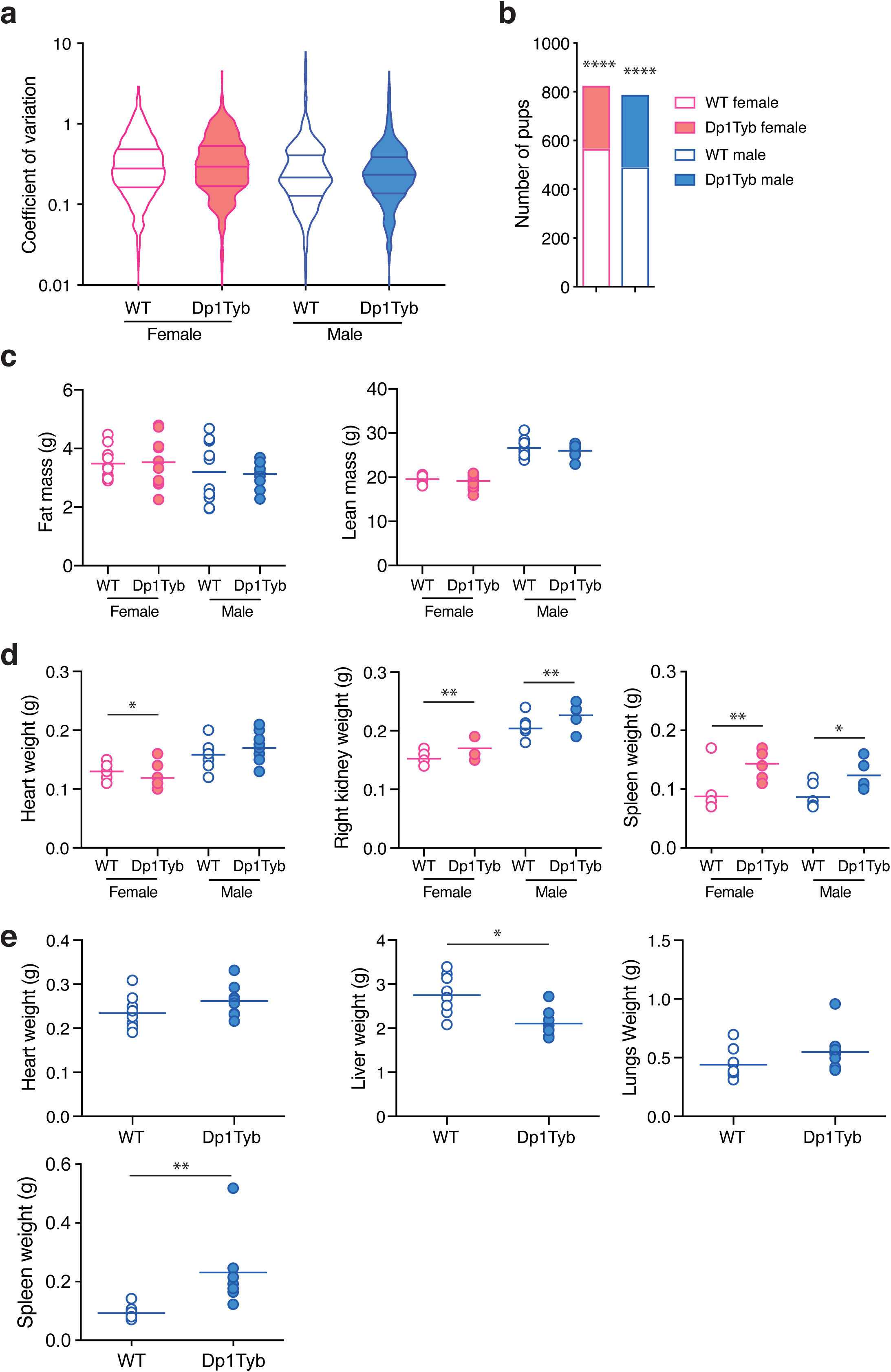
Decreased viability of Dp1Tyb mice. **a**, Coefficient of variation (standard deviation/mean) for all numerical parameters analysed shown as violin plots, with horizontal lines indicating median and 25^th^ and 75^th^ centiles. There are no significant differences between WT and Dp1Tyb measurements. Note the medians differ between females and males because some tests were carried out on only one sex (e.g. flow cytometric analysis was only carried out on females). **b**, Number of WT and Dp1Tyb mice recovered at weaning. **c**, Fat mass and lean mass of WT and Dp1Tyb mice (cohort 1) determined using ECHO-MRI. **d, e**, Weights of organs determined at 16 (d) and 57 (e) weeks of age. Horizontal lines indicate mean. * 0.01 < *q* < 0.05; ** 0.001 < *q* < 0.01; **** *q* < 0.0001.

**Supplementary Figure 2.**
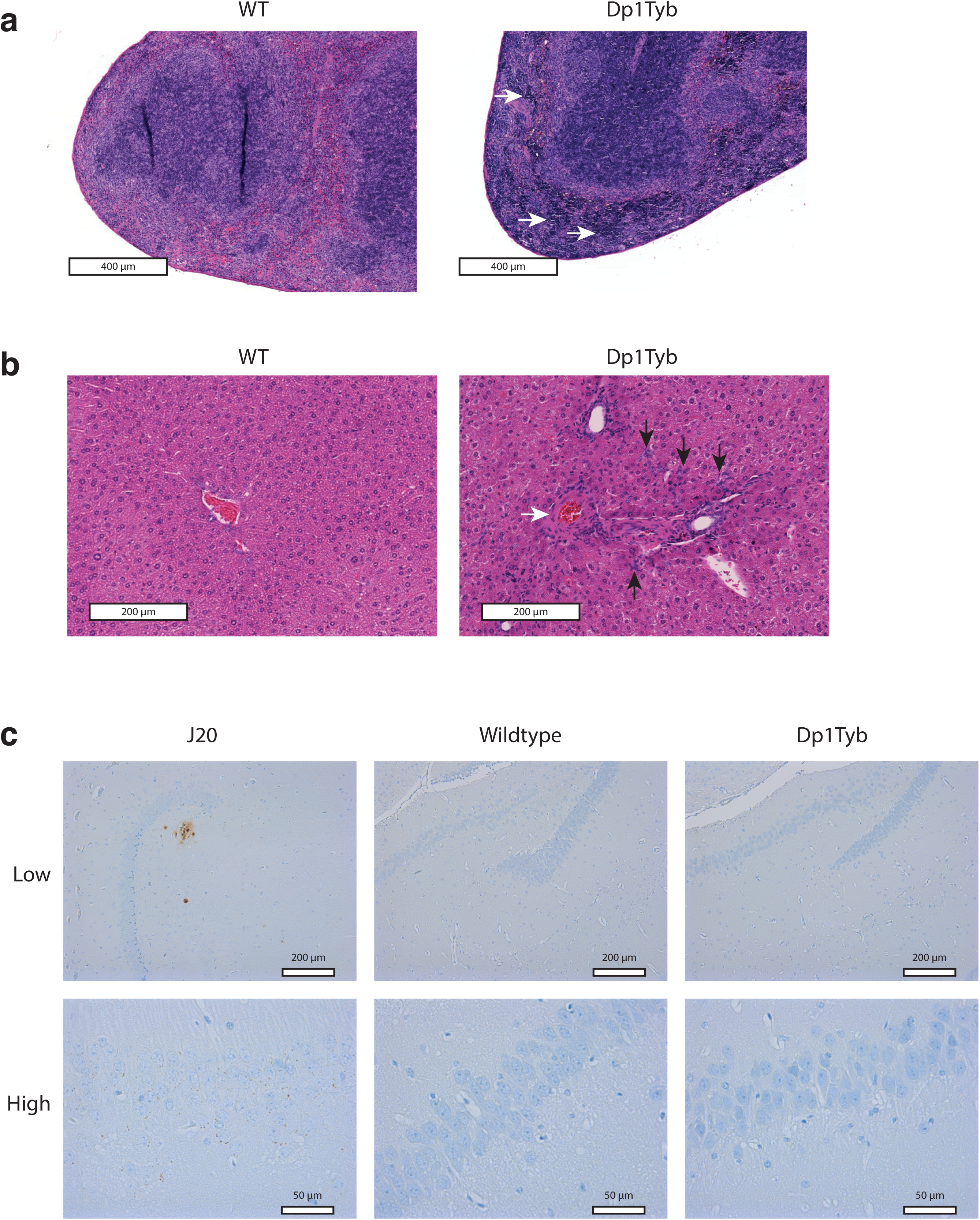
Pathological changes in spleens and livers of Dp1Tyb mice. **a**, Sections of spleens from WT and Dp1Tyb mice (cohort 1) stained with haematoxylin and eosin showing increased extramedullary haematopoiesis in Dp1Tyb mice (white arrows). **b**, Sections of livers from WT and Dp1Tyb mice (cohort 1) stained with haematoxylin and eosin showing prominent vessels (white arrow) and bile duct hyperplasia in Dp1Tyb mice (black arrows). **c**, Immunohistochemistry staining of Aβ-positive extracellular plaques (brown) in the CA3/dentate gyrus region of the hippocampus of a 42-week old J20 mouse used as a positive control, and in 57-week old WT and Dp1Tyb mice. Images shown at low (10x) and high (40x) magnification. Intracellular Aβ-positive staining was evident in the J20 CA3 pyramidal neurons only; both WT and Dp1Tyb mice were negative for intracellular CA3 neuronal Aβ staining.

**Supplementary Figure 3.**
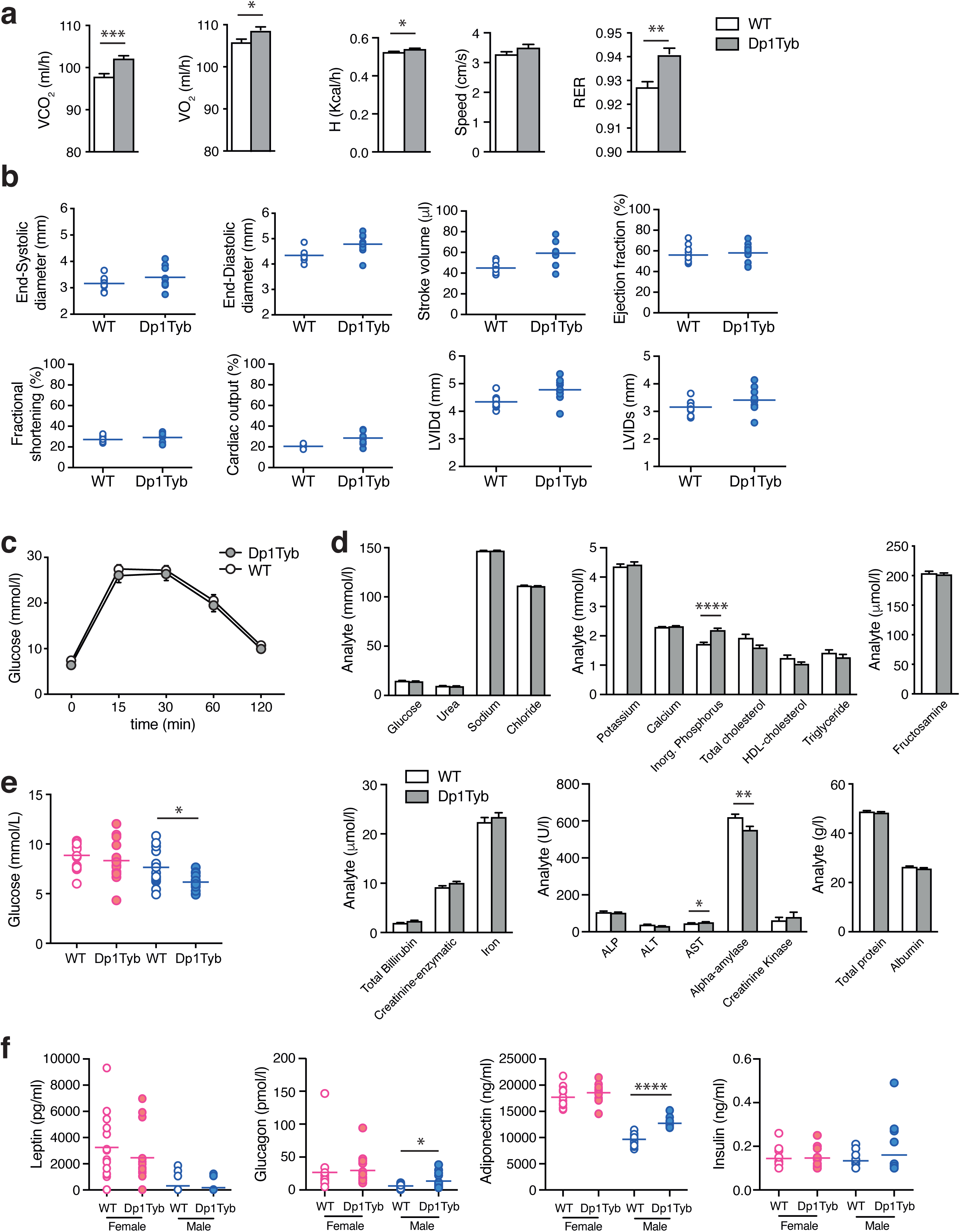
Calorimetry, ECG, glucose tolerance and plasma clinical chemistry of Dp1Tyb mice. **a**, Mean±SEM CO_2_ production (VCO_2_), O_2_ consumption (VO_2_), heat production (H), movement speed and respiratory exchange ratio (RER) of WT and Dp1Tyb mice (cohort 1). **b**, End-systolic diameter, end-diastolic diameter, stroke volume, ejection fraction, fractional shortening, cardiac output, left ventricular inner diameter in diastole (LVIDd) and left ventricular inner diameter in systole (LVIDs) in WT and Dp1Tyb mice at 57 weeks of age (cohort 5) determined by echocardiography. Horizontal lines indicate mean. **c**, Mean±SEM blood glucose level in WT and Dp1Tyb mice (cohort 1) injected with 20% glucose solution. **d**, Mean±SEM concentrations of the indicated analytes in the blood of free-fed WT and Dp1Tyb mice (cohort 1). **e**, Levels of glucose in fasted WT and Dp1Tyb mice (cohort 2). **f**, Levels of the indicated hormones in the blood of fasted WT and Dp1Tyb mice (cohort 2). Horizontal lines indicate mean. * 0.01 < *q* < 0.05; ** 0.001 < *q* < 0.01; *** 0.0001 < *q* < 0.001; **** *q* < 0.0001.

**Supplementary Figure 4.**
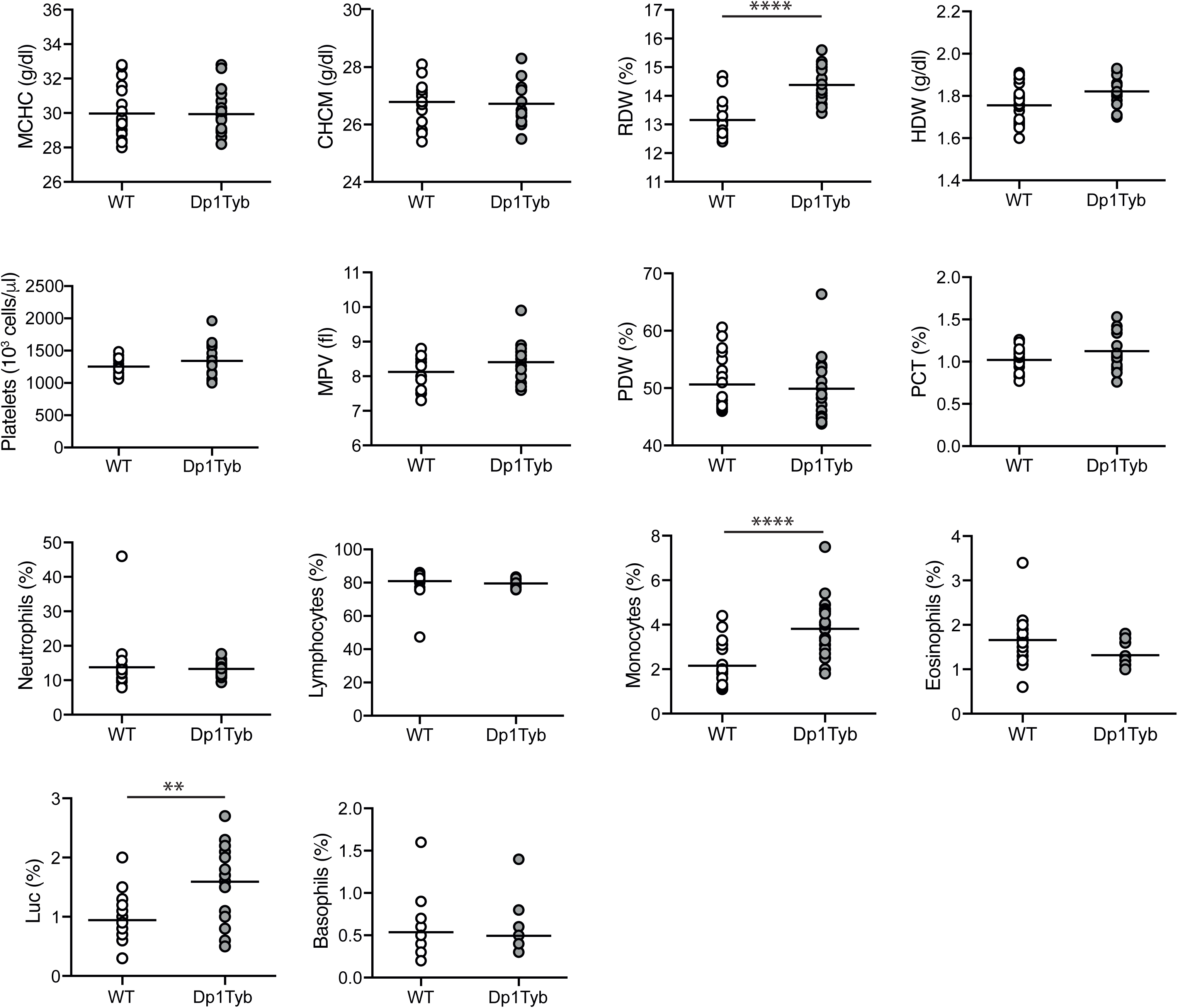
Haematology of Dp1Tyb mice. Mean corpuscular haemoglobin concentration (MCHC), cell haemoglobin concentration mean (CHCM), red blood cell distribution width (RDW), haemoglobin distribution width (HDW), platelet concentration, mean platelet volume (MPV), platelet distribution width (PDW), plateletcrit (PCT), percentage of neutrophils, lymphocytes, monocytes, eosinophils, large unstained cells (Luc) and basophils in the blood of WT and Dp1Tyb mice (cohort 1). ** 0.001 < *q* < 0.01; **** *q* < 0.0001.

**Supplementary Figure 5.**
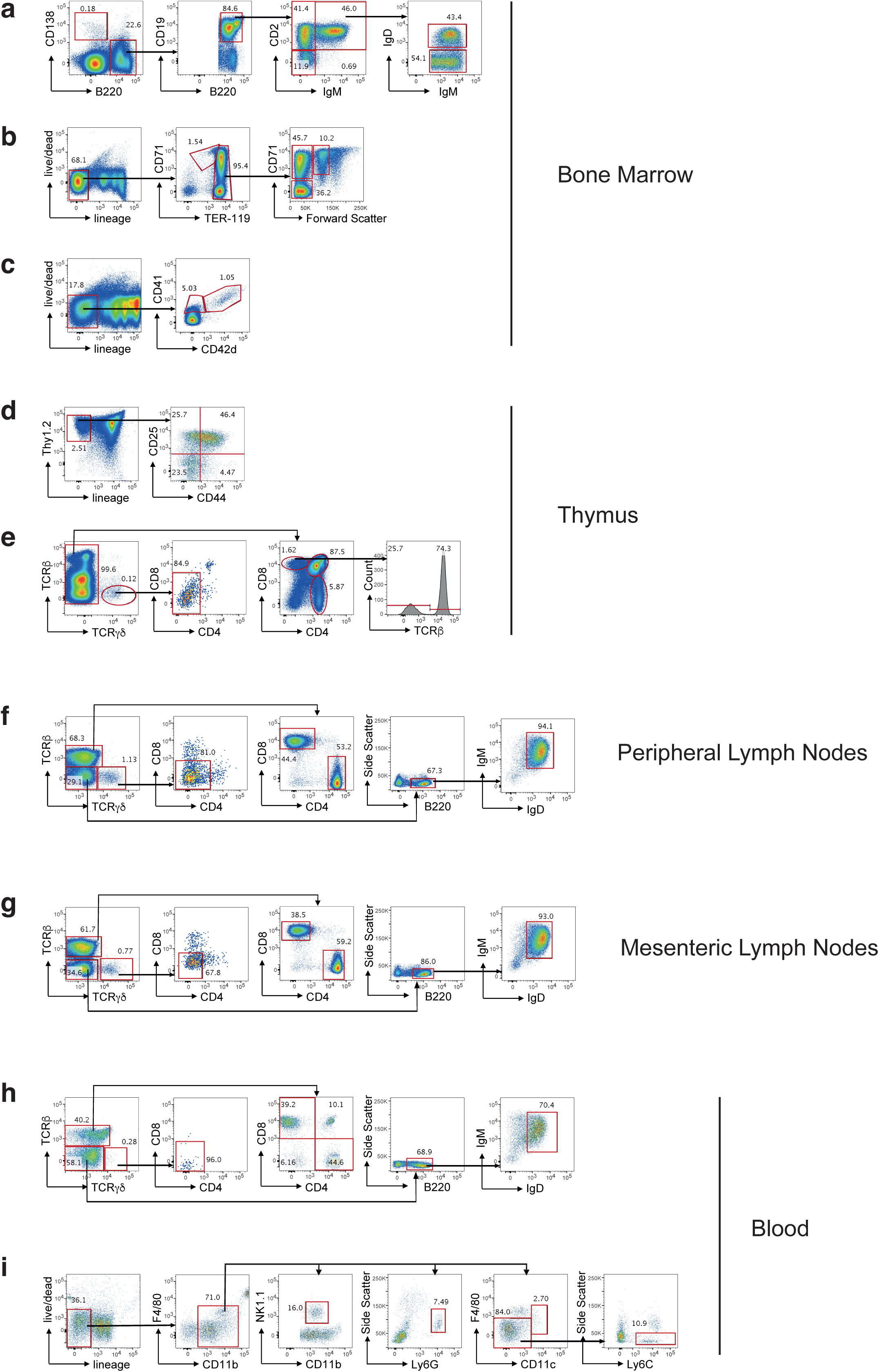
Flow cytometric gating strategies for analysis of bone marrow, thymus, lymph nodes, and blood. Cell types were identified using the following gating strategies. **a-c**, Bone marrow. **a**, plasma cells (B220^-^CD138^+^), pro-B (B220^+^CD2^-^IgM^-^), pre-B (B220^+^CD2^+^IgM^-^), immature (B220^+^CD2^+^IgM^+^IgD^-^), mature (B220^+^CD2^+^IgM^+^IgD^+^) B cells. **b**, Lineage^-^ cells (B220^-^CD3ε^-^Mac^-^1^-^Gr-1^-^) were subdivided into pro-erythroblasts (ProE, CD71^+^Ter119^lo^), EryA (Ter119^hi^CD71^+^FSC^hi^), EryB (Ter119^hi^CD71^+^FSC^lo^) and EryC (Ter119^hi^CD71^-^FSC^lo^) erythroid progenitors. **c**, immature (Lineage^-^CD41^+^CD42d^-^) and mature (Lineage^-^CD41^+^CD42d^+^) megakaryoblasts. **d, e,** Thymus. **d**, Double negative (DN) thymocytes defined as Lineage^-^ (CD4^-^CD8^-^TCRβ^-^TCRγδ^-^Gr-1^-^CD11b^-^CD11c^-^Dx5^-^Nk1.1^-^B220^-^) and Thy1.2^+^ were subdivided into DN1 (CD44^+^CD25^-^), DN2 (CD44^+^CD25^+^), DN3 (CD44^-^CD25^+^) and DN4 (CD44^-^CD25^-^). **e**, TCRγδ thymocytes (TCRβ^-^TCRγδ^+^CD4^-^CD8^-^), TCRγδ^-^ cells were subdivided into double positive (DP, CD4^+^CD8^+^), CD4^+^ single positives (SP, CD4^+^CD8^-^), CD8^+^ intermediate single positives (ISP, CD4^-^CD8^+^TCRβ^-^), and CD8^+^ SP (CD4^-^CD8^+^TCRβ^+^). **f-h,** Peripheral (f) and mesenteric (g) lymph nodes and blood (h). γδ T cells (TCRβ^-^TCRγδ^+^CD4^-^CD8^-^), TCRγδ^-^ cells were subdivided into CD4^+^ (TCRβ^+^CD4^+^CD8^-^) and CD8^+^ (TCRβ^+^CD4^-^CD8^+^) T cells, B cells (TCRβ^-^TCRγδ^-^ B220^+^IgM^+^IgD^+^). **i,** Blood. Lineage^-^ cells (B220^-^CD4^-^CD8^-^) were subdivided into NK cells (CD11b^lo^F4/80^-^NK1.1^+^), neutrophils (CD11b^lo^F4/80^-^Ly6G^+^SSC^hi^), dendritic cells (CD11b^lo^F4/80^-^CD11c^+^) and monocytes (CD11b^lo^F4/80^-^CD11c^-^Ly6C^+^). Numbers show percentage of cells falling into indicated gates.

**Supplementary Figure 6.**
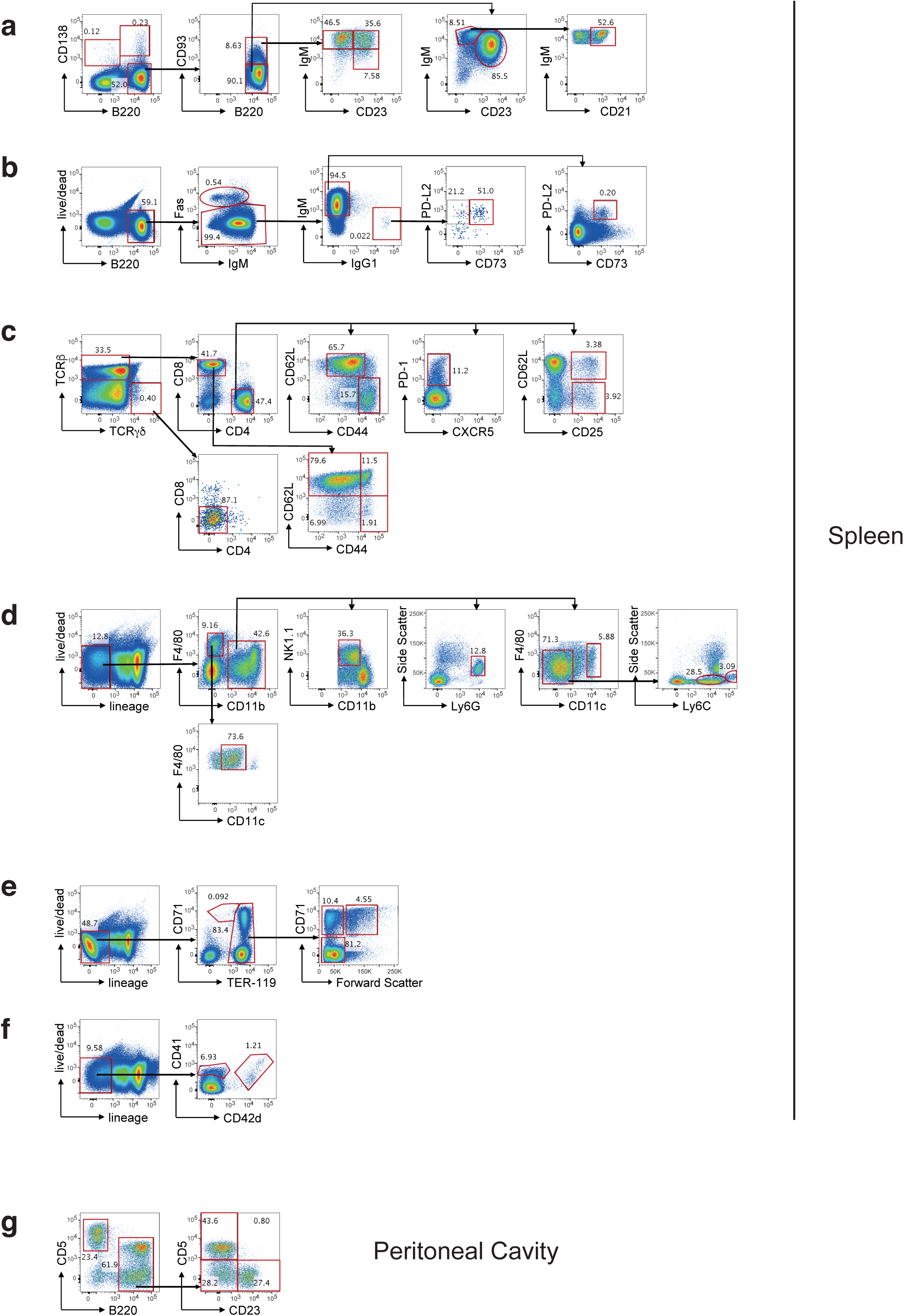
Flow cytometric gating strategies for analysis of spleen and peritoneal cavity. Cell types were identified using the following gating strategies. **a-f,** Spleen. **a**, plasmablasts (B220^+^CD138^+^), plasma cells (B220^-^CD138^+^), transitional B cells (B220^+^CD93^+^) were subdivided into type 1 (T1, IgM^+^CD23^-^), T2 (IgM^+^CD23^+^) and T3 (IgM^lo^CD23^+^), mature B cells (B220^+^CD93^-^) were subdivided into follicular (IgM^+^CD23^+^) and marginal zone (IgM^+^CD23^-^CD21^+^) B cells. **b**, Germinal centre (GC) B cells (B220^+^Fas^+^), IgM memory B cells (MBC) (B220^+^Fas^-^IgM^+^IgG1^-^CD73^+^PD-L2^+^) and IgG1 MBC (B220^+^Fas^-^IgM^-^IgG1^+^CD73^+^PD-L2^+^). **c**, γδ T cells (TCRβ^-^TCRγδ^+^CD4^-^ CD8^-^), TCRβ^+^TCRγδ^-^ cells were subdivided into total (CD4^+^), naïve (CD4^+^CD44^lo^CD62L^+^) and effector (CD4^+^CD44^hi^CD62L^-^) CD4^+^ T cells, T follicular helper T cells (CD4^+^PD-1^+^), regulatory T cells (CD4^+^CD25^+^CD62L^+ or -^), total (CD8^+^), naïve (CD8^+^CD44^-^CD62L^+^), central memory (CD8^+^CD44^+^CD62L^+^) and effector memory (CD8^+^CD44^+^CD62L^-^) CD8^+^ T cells. **d**, Lineage^-^ cells (B220^-^CD3ε^-^) were subdivided into macrophages (CD11b^-^CD11c^+^F4/80^+^), NK (CD11b^lo^F4/80^-^NK1.1^+^), neutrophils (CD11b^+^F4/80^-^Ly6G^+^SSC^hi^), dendritic cells (CD11b^+^F4/80^-^CD11c^+^) and monocytes (CD11b^+^F4/80^-^CD11c^-^) subdivided into Ly6C^lo^ and Ly6C^hi^ cells. **e**, Lineage^-^ cells (B220^-^CD3ε^-^Mac-1^-^Gr-1^-^) were subdivided into pro-erythroblasts (ProE, CD71^+^Ter119^lo^), EryA (Ter119^hi^CD71^+^FSC^hi^), EryB (Ter119^hi^CD71^+^FSC^lo^) and EryC (Ter119^hi^CD71^-^FSC^lo^) erythroid progenitors. **f**, immature (Lineage^-^CD41^+^CD42d^-^) and mature (Lineage^-^CD41^+^CD42d^+^) megakaryoblasts. **g**, Peritoneal cavity. B1a (B220^+^CD5^+^CD23^-^), B1b (B220^+^CD5^-^CD23^-^) and B2 (B220^+^CD5^-^CD23^+^) B cells and T cells (B220^-^CD5^+^). Numbers show percentage of cells falling into indicated gates.

**Supplementary Figure 7.**
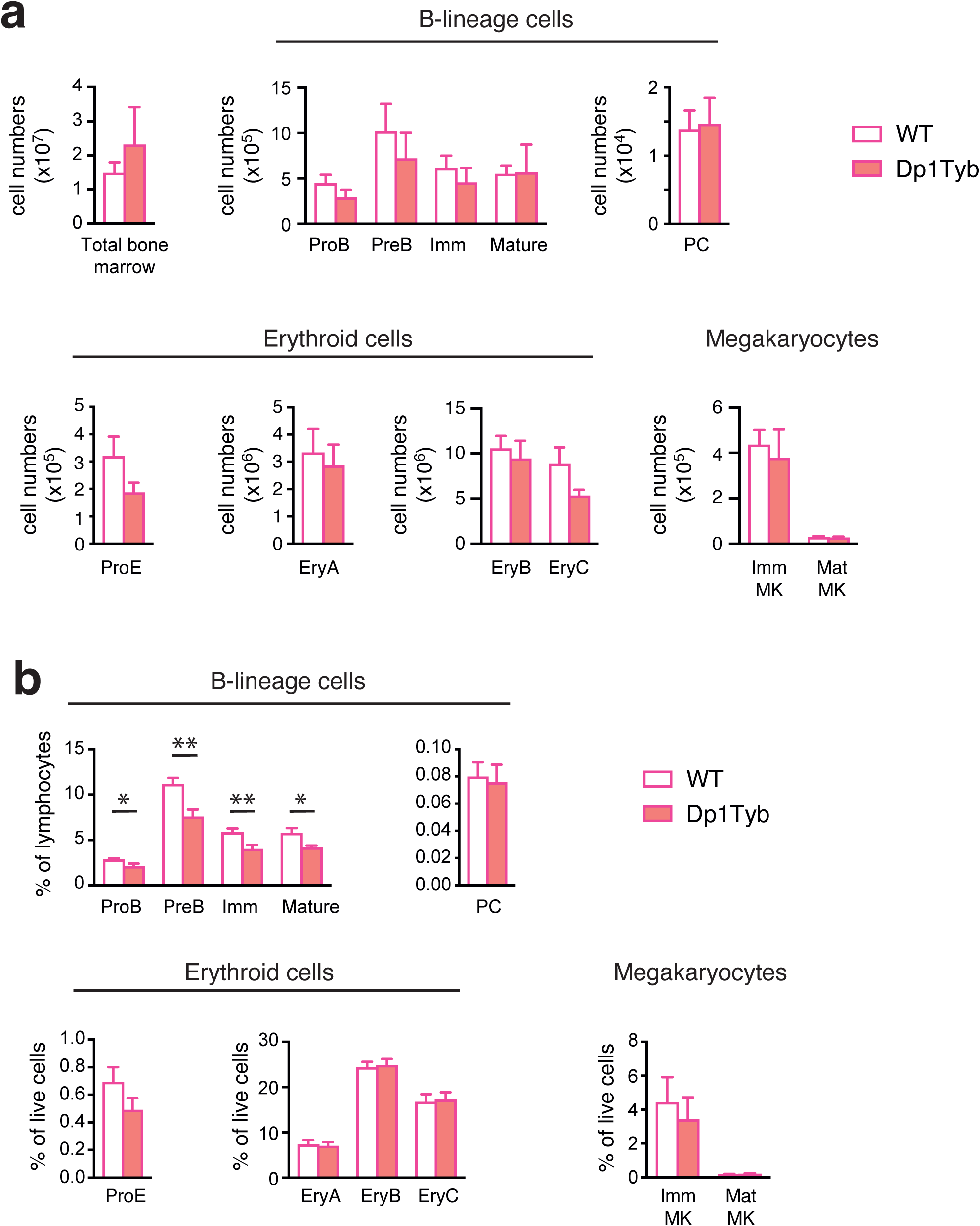
Flow cytometric analysis of bone marrow cells in Dp1Tyb mice. **a**, Mean±SEM number of cells in the bone marrow of WT and Dp1Tyb mice (cohort 4) (2 tibia and 2 femurs/mouse), showing total cells, pro-B, pre-B, immature (Imm) and mature B cells, plasma cells (PC), pro-erythroblasts (ProE), EryA, EryB and EryC erythroid progenitors, and immature (Imm) and mature (Mat) megakaryoblasts (MK). **b**, Mean±SEM percentages of the populations shown in a. * 0.01 < *q* < 0.05; ** 0.001 < *q* < 0.01.

**Supplementary Figure 8.**
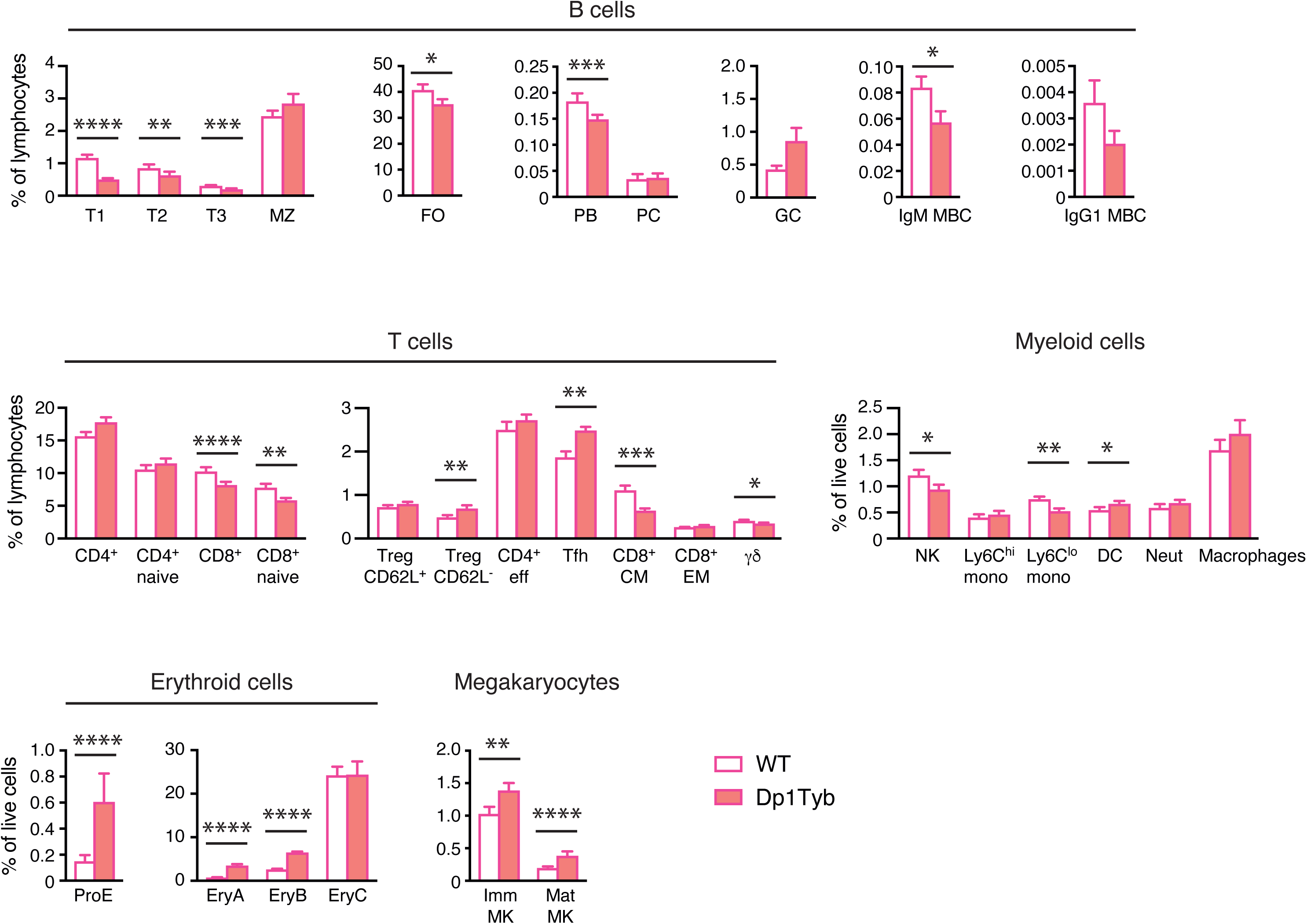
Flow cytometric analysis of splenocytes in Dp1Tyb mice. Mean±SEM percentages of transitional type 1 (T1), T2, T3, marginal zone (MZ), follicular (FO), germinal centre (GC) B cells, plasmablasts (PB), plasma cells (PC), IgM and IgG1 memory B cells (MBC), and total or naive CD4^+^ or CD8^+^ T cells, CD62L^+^ or CD62L^-^ regulatory T cells (Treg), CD4^+^ effector (eff) T cells, T follicular helper (Tfh) cells, CD8^+^ central memory (CM) and effector memory (EM) T cells, γδ T cells, NK cells, Ly6C^hi^ and Ly6C^lo^ monocytes (mono), dendritic cells (DC), neutrophils (Neut), macrophages, pro-erythroblasts (ProE), EryA, EryB and EryC erythroid progenitors, and immature (Imm) and mature (Mat) megakaryoblasts (MK) in the spleen of WT and Dp1Tyb mice (cohort 4). * 0.01 < *q* < 0.05; ** 0.001 < *q* < 0.01; *** 0.0001 < *q* < 0.001; **** *q* < 0.0001.

**Supplementary Figure 9.**
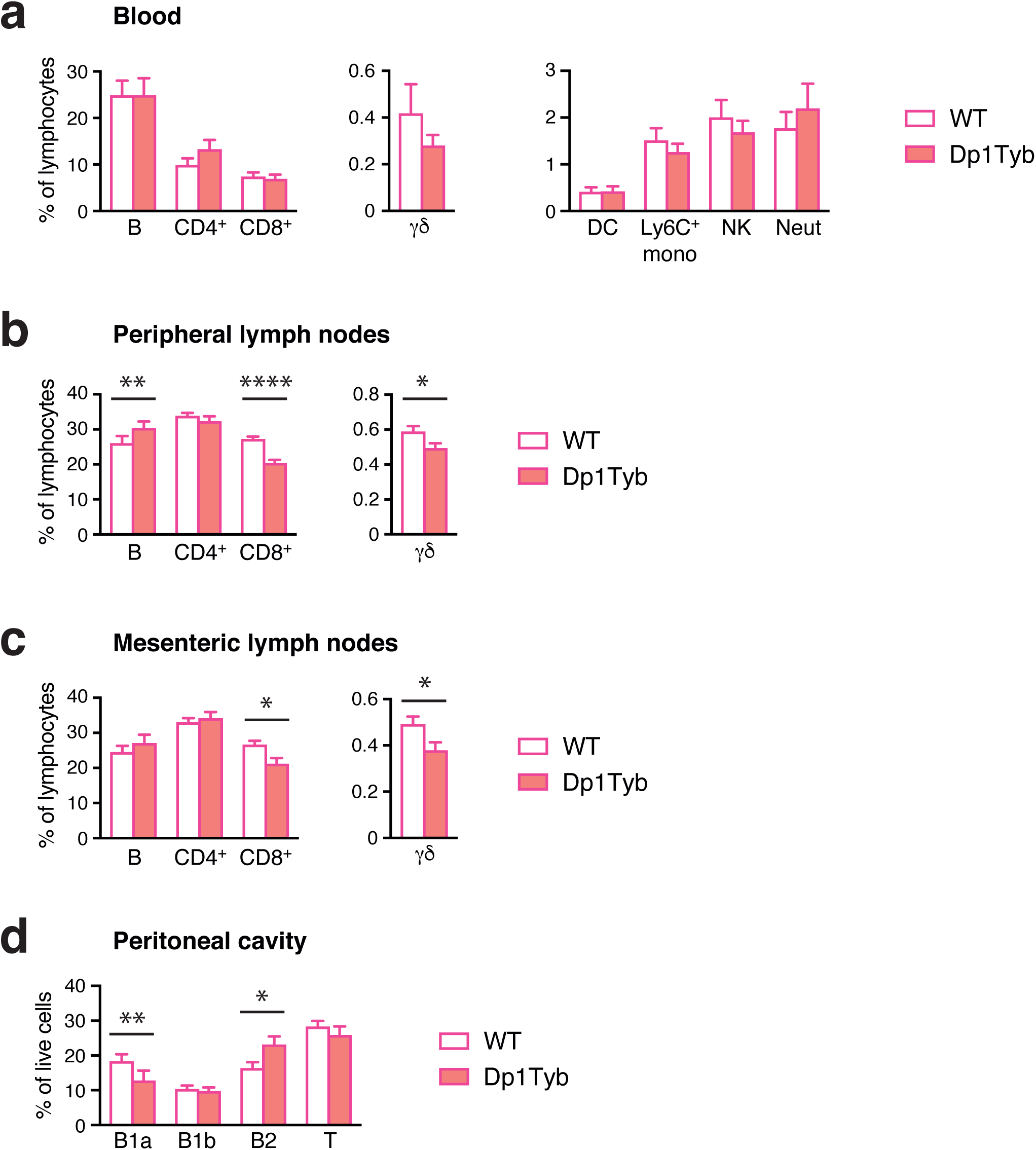
Flow cytometric analysis of blood, lymph nodes and peritoneal cavity in Dp1Tyb mice. **a-c**, Mean±SEM percentages of B cells, CD4^+^, CD8^+^ and γδ T cells, dendritic cells (DC), Ly6C^+^ monocytes, NK cells, and neutrophils (Neut) in the blood (a), peripheral lymph nodes (b), and mesenteric lymph nodes (c) of WT and Dp1Tyb mice (cohort 4). **d**, Mean±SEM percentages of B1a, B1b and B2 B cells and T cells in the peritoneal cavity of WT and Dp1Tyb mice (cohort 4). * 0.01 < *q* < 0.05; ** 0.001 < *q* < 0.01; **** *q* < 0.0001.

**Supplementary Figure 10.**
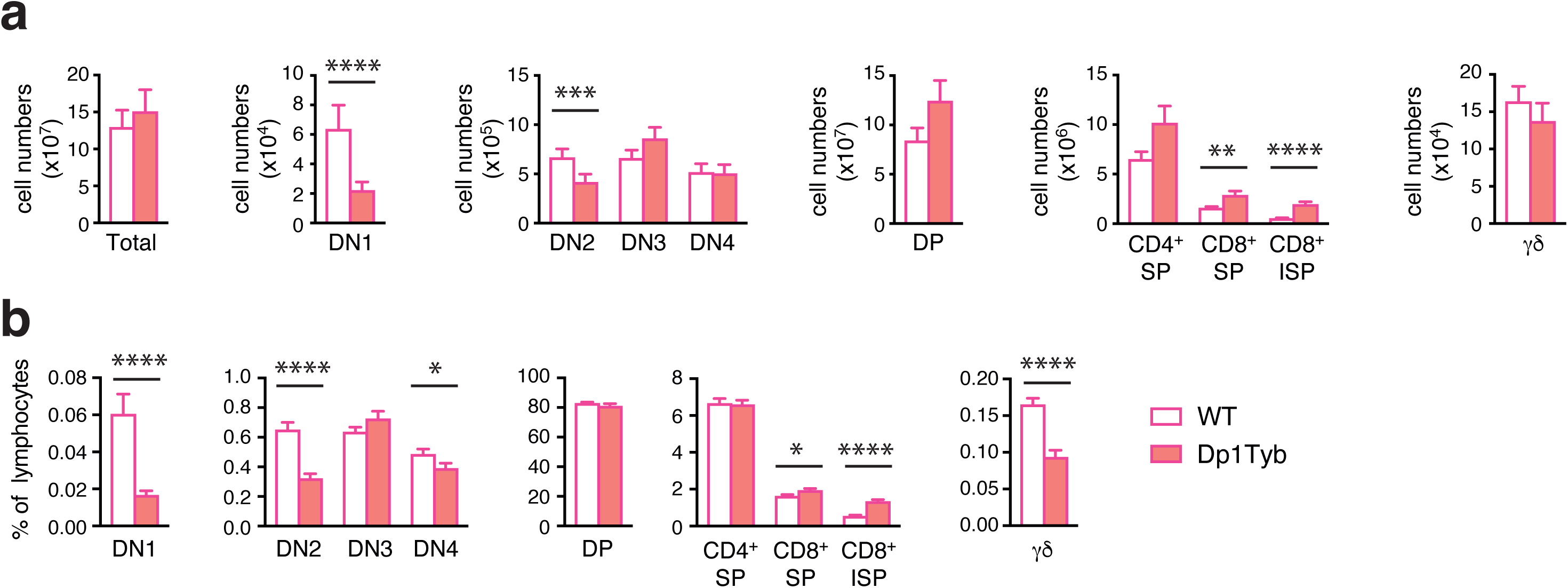
Flow cytometric analysis of thymocytes in Dp1Tyb mice. **a**, Mean±SEM number of cells in the thymus of WT and Dp1Tyb mice (cohort 4), showing total cells, DN1, DN2, DN3, DN4, DP, CD4^+^SP, CD8^+^SP, CD8^+^ISP and TCRγδ^+^ thymocytes. **b**, Mean±SEM percentages of the populations shown in a. * 0.01 < *q* < 0.05; ** 0.001 < *q* < 0.01; *** 0.0001 < *q* < 0.001; **** *q* < 0.0001.

**Supplementary Figure 11.**
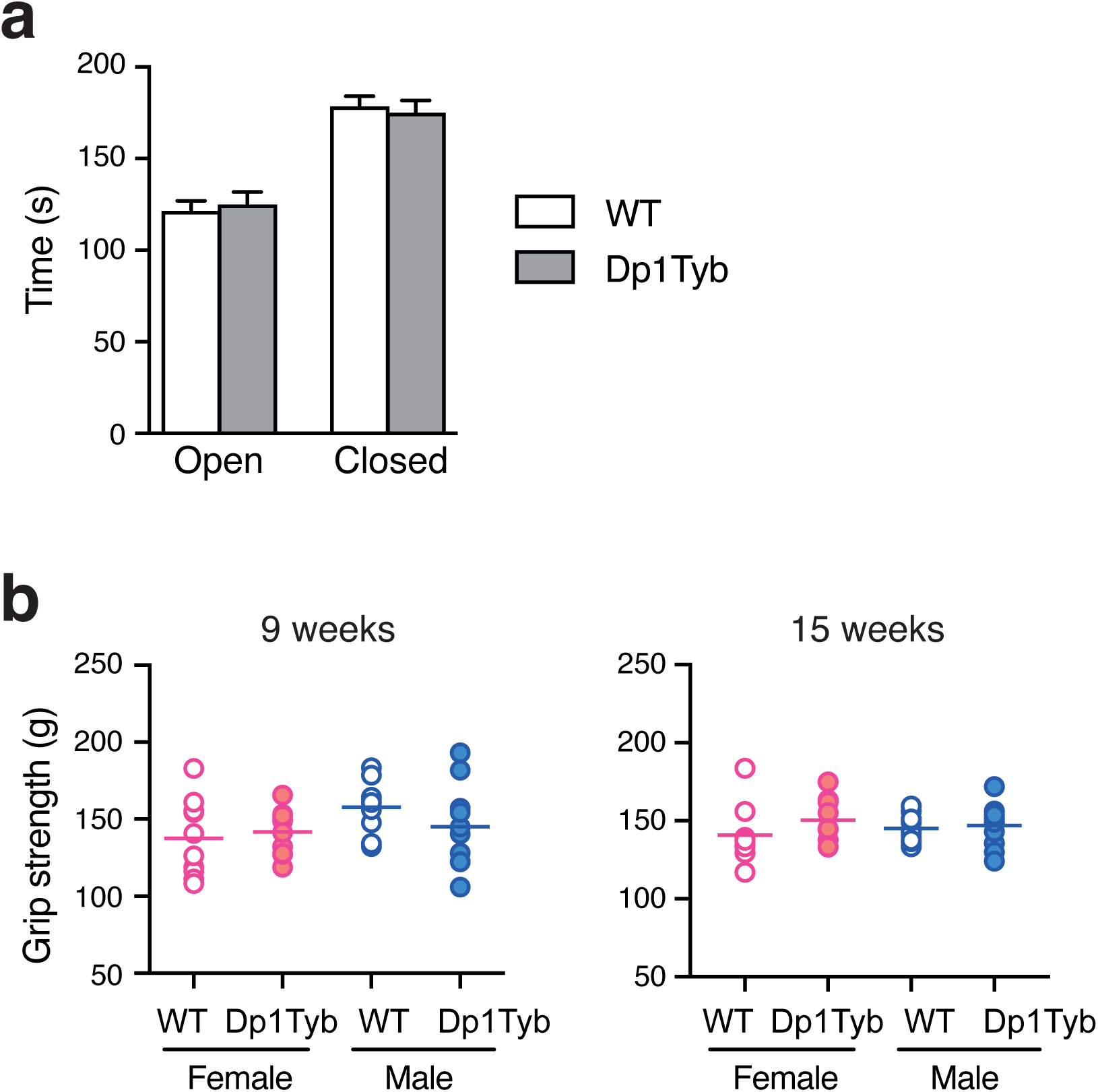
Open field, elevated zero maze and grip strength tests. **a**, Mean±SEM time spent in the open and closed arms of an elevated zero maze by WT and Dp1Tyb mice (cohort 3). **b**, Grip strength of WT and Dp1Tyb mice (cohort 1) determined using all four limbs at 9 and 15 weeks of age. Horizontal lines indicate mean.

